# Ablation catheter motion detection during contact force and VISITAG™ Module-guided pulmonary vein isolation

**DOI:** 10.1101/631374

**Authors:** David R. Tomlinson, Katie Biscombe, John True, Joanne Hosking, Adam J. Streeter

## Abstract

**Background:** During contact force (CF) and VISITAG™ Module (Biosense Webster) guided pulmonary vein isolation (PVI), ACCURESP™ respiratory motion adjustment is recommended, although without *in vivo* validation.

**Objective:** Since accurate LAPW radiofrequency (RF) annotation is crucial to avoid oesophageal thermal injury, we compared ACCURESP™ setting (“on” versus “off”) on RF annotation at the left atrial posterior wall (LAPW).

**Methods:** From a twenty-five patient cohort undergoing CF PVI (continuous RF, 30W) using general anaesthesia and VISITAG™ Module annotation-guidance (force-over-time 100% minimum 1g, 2mm position stability, ACCURESP™ “off”), respiratory motion detection occurred in eight, permitting retrospective comparison of ACCURESP™ settings.

*Results:* There were significant differences in LAPW RF data annotation according to ACCURESP™ setting. Comparing ACCURESP™ “on” versus “off”, respectively: Total annotated sites 82 versus 98; Median RF duration per-site 13.3s versus 10.6s (p<0.0001); Median force time integral 177g.s versus 130g.s (p=0.0002); Mean inter-lesion distance (ILD) 6.0mm versus 4.8mm (p=0.002). Considering only annotated site 1-to-2 transitions, three occurred with 0g CF; ACCURESP™ “on” minus “off” difference in RF duration was <0.6s. However, thirteen site 1-to-2 transitions during constant catheter-tissue contact (ILD range 2.1 – 7.0mm) demonstrated a mean difference in annotated RF duration at site 1, of 3.7s (Range: −1.3 – 11.3s). Reconstituted curves displaying catheter position data, CF, impedance and site 1-to-2 transition according to ACCURESP™ setting, demonstrated multiple markers of catheter movement coinciding with ACCURESP™ “off”.

*Conclusions:* ACCURESP™ “off” setting demonstrated excellent catheter movement detection properties and represents an optimal method towards the annotation of stable sites of RF delivery at the LAPW.

## Introduction

The central importance of electrical isolation of the pulmonary veins (PVs) towards eliminating atrial fibrillation (AF) is well-established^1,2^, but achieving this end-point using catheter ablation involves three principal challenges: (1) There is no universally applicable and validated means to directly visualise ablation lesion creation, either in terms of the transmural (TM) extent or radius; (2) The left atrial (LA) wall thickness is variable^3^, yet unknown to the operator, and while partial wall-thickness lesions result in electrical conduction gaps and recurrent AF^2,4^, excessive energy delivery risks life-threatening extra-cardiac thermal trauma^5^, and; (3) Cardiac and respiratory cycle-induced motion creates a moving target for radiofrequency (RF)-based catheter ablation technologies.

Theoretically, idealised pulmonary vein isolation (PVI) protocols are likely to involve variable energy dosing according to the LA wall thickness, with real-time incorporation of suitable measures of the tissue effects of energy delivery – i.e. lesion transmurality and radius – into a 3D “model” using objective and validated methods. When supported by suitable methodology for defining a stable point of catheter-tissue interaction during RF application, such “modelling” should facilitate the knowing completion of permanent PVI, without extra-cardiac thermal trauma, in all but exceptional cases and for all suitably skilled operators.

Important progress towards this goal has been reported in studies utilising the objective RF annotation module VISITAG™ (Biosense Webster Inc., Diamond Bar, CA), with a coordinated series of methodological steps towards tailored contact force (CF)-guided PVI lesion sets, including targeting minimum permissible inter-lesion distance (ILD) and site-specific RF energy delivery (i.e. anterior versus posterior wall) according to a weighted formula incorporating measures of RF power, duration and CF.^6,7^ However, although very high clinical success has been achieved in selected operators’ practice, these protocols fail to represent perfect descriptors of reproducible and suitably tailored PVI protocols in four important respects:

(1) Studies of RF delivery during PVI in humans have provided evidence of greater effect at left-sided LA posterior wall (LAPW) sites^8,9^, therefore any protocol without suitable energy dosing adjustment is likely to incur increased risk of extra-cardiac thermal trauma;

(2) There was no incorporation of measures of the TM tissue response to RF delivery, yet *in vivo* animal studies have demonstrated that a change in the unipolar electrogram (UE) morphology from RS to “pure R” may be indicative of histologically-confirmed TM lesions.^10,11^ Furthermore, a study conducted in humans has confirmed the utility of RF titration according to real-time assessments of pure R UE morphology change towards a highly effective CF-guided PVI protocol^12^;

(3) The chosen VISITAG™ Module CF filter settings (force-over-time 30%, minimum 5g^13^ and 30% minimum 4g^7^) permit variable out-of-phase catheter-tissue interaction due to intermittent catheter-tissue contact (i.e. 0g CF), though with RF annotation on-going and displayed / modelled as a single point;

(4) Employing either conscious sedation or general anaesthesia (GA) with intermittent positive pressure ventilation (IPPV), these studies^13^ routinely utilised ACCURESP™ respiratory adjustment (Biosense Webster) “on” for VISITAG™ Module annotation (Molloy Das, personal communication). However, ACCURESP™ remains without *in vivo* validation as a component of automated RF annotation and lesion modelling methodology.

Therefore, the purpose of this present report was to retrospectively investigate the effects of ACCURESP™ respiratory adjustment setting on RF lesion modelling using the VISITAG™ Module, thereby determining the most appropriate ACCURESP™ setting for RF lesion annotation during CF and VISITAG™ Module-guided PVI.

## Methods

CF and VISITAG™ Module-guided PVI was performed by a single-operator employing a previously reported standardised protocol^9^ in a consecutive series of unselected adult patients with symptomatic AF undergoing first-time PVI according to current treatment indications.^14^ Briefly, all procedures were undertaken using GA and IPPV, with ACCURESP™ respiratory training undertaken pre-ablation and applied as required to complete the CARTO^®^3 geometry (V.3, Biosense Webster). Specifically, ACCURESP™ training was first performed with a LASSO^®^Nav catheter (Biosense Webster, 2-5-2mm inter-electrode spacing) placed in the right superior PV. If there was insufficient respiratory motion to trigger the ACCURESP™ detection threshold, the catheter was placed in the left inferior PV and ACCURESP™ training re-checked. The tidal volume was never deliberately increased, so ACCURESP™ respiratory motion detection was negative in some cases. Importantly, even in cases where the ACCURESP™ respiratory adjustment threshold was triggered, the ACCURESP™ setting “on” was never prospectively applied to the VISITAG™ Module filter preferences during ablation.

Temperature-controlled RF at 30W (17ml/min irrigation) was delivered via a ThermoCool® SmartTouch® catheter using Agilis™ NxT sheath (Abbott, St Paul, MN) support during proximal pole CS pacing at 600ms. VISITAG™ Module filter preferences for automated RF annotation were: Positional stability range 2mm, tag display duration 3s; force-over-time 100% minimum 1g (the latter derived from a previous study^15^ and designed to ensure on-going RF annotation only in the presence of constant catheter-tissue contact). Lesion placement was guided by VISITAG™ Module annotation, with the preferred site of first RF application at the LAPW opposite each superior PV ∼1cm from the PV ostium; in cases where constant catheter-tissue contact could only be achieved with maximal CF ≥70g, an adjacent LAPW site with lower peak CF was chosen. The target annotated RF duration at each first-ablated LAPW site, as well as any subsequent “RF ON” sites and at the carina (if ablated) was 15s, whereas ∼9-11s was the target for all other sites consecutively annotated during continuous RF delivery. Following the required period of first-site annotated RF, target ILD ≤6mm was achieved predominantly during continuous RF application using rapid movement of the catheter tip initiated via the Agilis sheath, aided by the distance measurement tool; point-by-point RF was also applied as necessary. Following completion of circumferential PVI (entrance and exit block), spontaneous recovery of PV conduction was assessed and eliminated during a minimum 20-minute wait; dormant recovery was evaluated and eliminated a minimum of 20 minutes after the last RF. Neither oesophageal luminal temperature monitoring nor post-ablation endoscopic evaluation was employed.

For all cases where ACCURESP™ triggering threshold was exceeded, VISITAG™ Module annotated RF and UE morphology change data were retrospectively collected. The focus for analysis was all ablation-naïve LAPW sites (i.e. first encirclement, not including touch-up lesions), since ablation here entails risk of atrio-oesophageal fistula (AEF), therefore accurate lesion modelling via automated annotation methodology is of particular importance in this region. Annotated RF duration, mean CF, force time integral (FTI) and impedance drop data for each site were obtained via the VISITAG™ Module export function. To examine the effects of ACCURESP™ setting, data export was performed separately for ACCURESP™ “on and “off”. ILD was determined on-line using the proprietary measurement tool. Retrospective UE analysis was performed via CARTOREPLAY™ (Biosense Webster) as previously described^9^; electrograms are automatically deleted at 12-18 hours after case completion (a CARTO^®^3 system function), so UE morphology data was only obtained for the ACCURESP™ “off” setting. This work received IRB approval for publication as a retrospective service evaluation. All patients provided written, informed consent.

### Analysis

The question was whether ACCURESP™ “on” versus “off” settings resulted in significant differences in the number of annotated LAPW sites, annotated RF duration, ILD, total impedance drop, mean CF and FTI. ACCURESP™ “on” versus “off” annotation timing accuracy was assessed at all site 1-to-2 ablation catheter transitions (i.e. site 1-to-2 transitions at the commencement of each of the left and right PVI encirclement) by visual inspection of reconstituted catheter position, CF and impedance curves. These were considered important since the accuracy of ablation catheter motion detection away from any annotated site of RF delivery is equally important for both point-by-point and continuous RF ablation protocols. Ablation catheter motion associated with transition between annotated sites 1-to-2 was identified according to either of the following criteria:

(1) Site 1 annotation “end” due to a 0g CF event (i.e. CF breaching the VISITAG™ Module force-over-time filter of 100% minimum 1g);

(2) Inter-ablation site transition during constant catheter-tissue contact, but accompanied by UE morphology change from pure R at site 1 completion, to RS at site 2 onset (i.e. indicating movement from a site of TM ablation effect to an adjacent ablation naive site).

By definition, all remaining site 1-to-2 transitions occurred during constant catheter-tissue contact and were effected with pure R UE morphology at both site 1 completion and site 2 onset; subsequent LAPW annotated transitions (sites 2-to-3, 3-to-4 and 4-to-5) were similarly identified and described.

VISITAG™ Module data comprising the x,y,z coordinates of the catheter tip position unadjusted for respiratory motion (i.e. using the “RawPositions” data file) and measured at 60 positions per second were exported to R^16^ for analysis. The standard deviation (SD) over a rolling one second period – as utilised by the CARTO^®^3 system for position stability annotation – was calculated in R from the Euclidean distances shifted between each position. These “position stability” data were plotted according to the patient interface unit (PIU) “system” time and supplemented with the absolute CF (at 20Hz) and impedance (at 10Hz) data derived from the exported “ContactForceData” “AblationData” files respectively, and end-expiration timing data from the “EndExperium” file. Finally, PIU start and stop times for all annotated sites according to each ACCURESP™ setting were obtained from the respective “AblationSites” files.

### Statistics

Exported text files were converted to Excel data and imported to GraphPad Prism version 4.03 (GraphPad Software, San Diego, CA) for analysis. Comparisons were made between the ACCURESP™ “on” and “off” settings based on the means and standard deviations of biophysical data (i.e. ILD, RF duration, CF, impedance drop and FTI), or medians along with the 1^st^ and 3^rd^ quartile (IQR), where these were determined to be skewed. Differences in the biophysical data between the ACCURESP™ “on” versus “off” settings were tested using the unpaired t-test, or the Mann-Whitney test where data could not be assumed to be normally distributed. The Pearson correlation coefficient was calculated to determine the strength of association between ACCURESP™ set “off” ILD and the difference (ACCURESP™ “on” minus “off”) in RF duration, FTI and ILD. In this exploratory analysis, significance was set at the 5% level.

## Results

Twenty-five patients underwent first-time PVI as described, between November 2016 and May 2017: 13 persistent AF, 12 PAF; 19 male (76%); mean age 57 [SD: 14] years and mean CHA_2_DS_2_-VASc score 1.3 [SD: 1.3]. Complete PVI was achieved in all without spontaneous / dormant recovery of PV conduction, following mean 16.2 [SD: 3.1] minutes of RF, with no procedural complications. The ACCURESP™-triggering cohort comprised 8 of 25 cases (32%); considering age, body mass index and RF duration required for case completion, there were no significant differences between the cohorts with and without ACCURESP™ threshold triggering.

Comparing ACCURESP™ “on” versus “off”, the number of annotated LAPW sites and total LAPW RF duration were 82 and 98, and 1091s and 1006s, respectively (annotated biophysical RF data according to ACCURESP™ setting are shown in table 1). For each group of annotated sites (i.e. left or right-sided), per-site RF duration and FTI were significantly greater with ACCURESP™ “on” versus “off”; i.e. mean left-sided RF duration 13.1s versus 9.9s (p=0.0003) and median FTI 156g.s versus 114g.s (p=0.0003), respectively and mean right-sided RF duration 13.5s versus 10.6s (p=0.006) and median FTI 228g.s versus 166g.s (p=0.04), respectively. Analysis of the combined left and right sides also demonstrated significantly greater mean ILD with ACCURESP™ “on”; i.e. 6.0mm versus 4.8mm (p=0.002).

**Table 1:** Biophysical data at all annotated sites of RF delivery at the LAPW according to ACCURESP™ setting (i.e. “on” versus “off”) and site; data shown are mean [SD] or median (1^st^ – 3^rd^ quartile), as appropriate. Note: “Number of annotated sites” data represents the mean of 8 patients; ^†^indicates the number of LAPW annotated sites identified and used for analyses of the remaining variables.

|  | Left-sided annotated sites |  |  | Right-sided annotated sites |  |  | All annotated sites |  |  |
| --- | --- | --- | --- | --- | --- | --- | --- | --- | --- |
|  | ACCURESP<br>ON (N=43<sup>†</sup>) | ACCURESP<br>OFF (N=52<sup>†</sup>) | *p* | ACCURESP<br>ON (N=39<sup>†</sup>) | ACCURESP<br>OFF (N=46<sup>†</sup>) | *p* | ACCURESP<br>ON (N=82<sup>†</sup>) | ACCURESP<br>OFF (N=98<sup>†</sup>) | *p* |
| Number of sites<br>per patient | 5.4<br>[1.3] | 6.5<br>[1.6] | 0.15 | 4.9<br>[1.5] | 5.8<br>[1.7] | 0.28 | 5.1<br>[1.4] | 6.1<br>[1.6] | 0.07 |
| ILD (mm) | 5.6<br>[1.6] | 4.6<br>[1.7] | 0.01 | 6.2<br>[2.2] | 5.4<br>[2.6] | 0.16 | 6.0<br>(4.7 – 6.8) | 4.8<br>(3.9 – 5.6) | 0.002 |
| RF duration (s) | 13.1<br>[4.4] | 9.9<br>[3.9] | 0.0003 | 13.5<br>[5.5] | 10.6<br>[3.9] | 0.006 | 13.3<br>(10.3 – 15.6) | 10.6<br>(7.5 – 13.2) | <0.0001 |
| Mean CF (g) | 12.1<br>(9.9 – 15.6) | 11.5<br>(9.7 – 14.9) | 0.67 | 17.9<br>(13.4 – 21.6) | 18.6<br>(12.1 – 22.1) | 0.68 | 14.4<br>(10.8 – 19.7) | 14.0<br>(10.7 – 19.9) | 0.85 |
| Impedance drop<br>(Ω) | 8.0<br>(4.3 – 12.2) | 6.6<br>(3.4 – 10.6) | 0.24 | 5.9<br>(1.5 – 8.8) | 3.7<br>(0.8 – 7.1) | 0.34 | 6.5<br>(3.5 – 10.0) | 5.4<br>(1.7 – 8.9) | 0.14 |
| FTI (g.s) | 156<br>(123 – 204) | 114<br>(31 – 161) | 0.0003 | 228<br>(134 – 315) | 166<br>(111 – 243) | 0.04 | 177<br>(133 – 250) | 130<br>(104 – 195) | 0.0002 |

A comparison of annotated biophysical data at annotated site 1 according to ACCURESP™ setting is shown in table 2: At the left-side and comparing ACCURESP™ “on” versus “off” the observed distance to site 2 (6.6mm versus 5.3mm, p=0.07), RF duration (16.0s versus 15.1s, p=0.16) and FTI (185g.s versus 163g.s, p=0.33) were greater with ACCURESP™ “on” but the differences were not statistically significant. At the right-side the observed distance to site 2 (7.2mm versus 5.0mm, p=0.13) was greater with ACCURESP™ “on” but the difference was not statistically significant. Combined data analysis demonstrated that annotated distance to site 2 was significantly greater with ACCURESP™ “on” versus “off” (i.e. 6.7mm versus 5.2mm, p=0.02), while the observed difference in site 1 RF duration (15.7s versus 15.1s, p=0.09) and FTI (240g.s versus 198g.s, p=0.38) was not statistically significant.

**Table 2:** Annotated biophysical data at first-annotated LAPW sites according to ACCURESP™ setting (i.e. “on” versus “off”) and site; data shown are median (1^st^ – 3^rd^ quartile).

|  | Left-sided 1<sup>st</sup> annotated site |  |  | Right-sided 1<sup>st</sup> annotated site |  |  | All 1<sup>st</sup> annotated sites |  |  |
| --- | --- | --- | --- | --- | --- | --- | --- | --- | --- |
|  | ACCURESP<br>ON (N=8) | ACCURESP<br>OFF (N=8) | *p* | ACCURESP<br>ON (N=8) | ACCURESP<br>OFF (N=8) | *p* | ACCURESP<br>ON (N=16) | ACCURESP<br>OFF (N=16) | *p* |
| Distance to site 2<br>(mm) | 6.6<br>(6.3 – 7.0) | 5.3<br>(4.6 – 6.3) | 0.07 | 7.2<br>(6.3 – 8.4) | 5.0<br>(4.5 – 8.1) | 0.13 | 6.7<br>(6.3 – 7.8) | 5.2<br>(4.5 – 6.8) | 0.02 |
| RF duration (s) | 16.0<br>(15.2 – 18.7) | 15.1<br>(14.7 – 15.6) | 0.16 | 15.5<br>(14.7 – 23.7) | 15.1<br>(14.7 – 15.7) | 0.57 | 15.7<br>(14.9 – 22.2) | 15.1<br>(14.7 – 15.6) | 0.09 |
| Mean CF (g) | 10.8<br>(9.5 – 12.9) | 10.8<br>(9.0 – 13.0) | 0.88 | 16.8<br>(12.6 – 21.6) | 17.8<br>(11.7 – 21.2) | 1.00 | 12.9<br>(10.1 – 18.2) | 13.0<br>(9.0 – 18.3) | 0.84 |
| Impedance drop<br>(Ω) | 13.5<br>(11.5 – 23.6) | 13.5<br>(10.9 – 21.5) | 0.80 | 10.3<br>(8.5 – 12.4) | 10.3<br>(7.8 – 12.4) | 0.96 | 12.0<br>(9.1 – 15.7) | 11.5<br>(9.1 – 15.2) | 0.81 |
| FTI (gs) | 185<br>(150 - 230) | 163<br>(123 - 198) | 0.33 | 268<br>(240 - 376) | 264<br>(185 - 327) | 0.65 | 240<br>(163 - 322) | 198<br>(135 - 270) | 0.38 |

### Analyses at sites of deliberate catheter movement: Site 1-to-2 annotated transition

Comparing the annotated biophysical data (ACCURESP™ “on” minus “off”) at three site 1-to-2 annotated transitions concurrent with 0g CF, the maximum difference in annotated RF duration, FTI and ILD was −0.6s, −17g.s and 2.2mm respectively, with no difference in annotated impedance drop (table 3 and figures 1-3).

**Table 3:** Annotated biophysical data at site 1-to-2 transition, with deliberate catheter movement identified according to criteria of a 0g CF event, or change in the UE morphology from pure R (site 1 completion) to RS (site 2 onset). Annotated site 1 RF duration, site 1-to-2 ILD, impedance drop and FTI data are displayed according to ACCURESP™ setting, with the difference (“Diff” – i.e. ACCURESP™ “on” minus “off”) also shown; PV, pulmonary vein.

| Case number | PV | Position instability criterion | Site 1 RF duration (s) | | | Distance to site 2 (mm) | | | Site 1 impedance drop ($\Omega$) | | | Site 1 FTI (g.s) | | |
| --- | --- | --- | --- | --- | --- | --- | --- | --- | --- | --- | --- | --- | --- | --- |
|  |  |  | ACCURESP |  |  | ACCURESP |  |  | ACCURESP |  |  | ACCURESP |  |  |
|  |  |  | ON | OFF | *Diff* | ON | OFF | *Diff* | ON | OFF | *Diff* | ON | OFF | *Diff* |
| 5 | Right | 0g CF | 15.5 | 15.6 | -0.1 | 6.3 | 4.1 | 2.2 | 9.4 | 9.4 | 0 | 228 | 228 | 0 |
| 10 | Right | 0g CF | 14.4 | 14.4 | 0 | 8.6 | 7.8 | 0.8 | 25.3 | 25.3 | 0 | 261 | 261 | 0 |
| 11 | Right | 0g CF | 14.0 | 14.6 | -0.6 | 8.6 | 6.5 | 2.1 | 6.5 | 6.5 | 0 | 382 | 399 | -17 |
| 5 | Left | Pure R UE to RS | 26.5 | 15.2 | 11.3 | 6.7 | 4.1 | 2.6 | 29.5 | 26.2 | 3.3 | 256 | 117 | 139 |
| 11 | Left | Pure R UE to RS | 20.9 | 18.6 | 2.3 | 4.9 | 5.2 | -0.3 | 12.2 | 11.1 | 1.1 | 482 | 434 | 48 |
| 22 | Right | Pure R UE to RS | 15.1 | 14.8 | 0.3 | 6.1 | 5.0 | 1.1 | 11.1 | 11.1 | 0 | 274 | 272 | 2 |
| 23 | Right | Pure R UE to RS | 15.5 | 15.0 | 0.5 | 8.1 | 8.8 | -0.7 | 11.9 | 11.9 | 0 | 120 | 112 | 8 |

**Figure 1:**
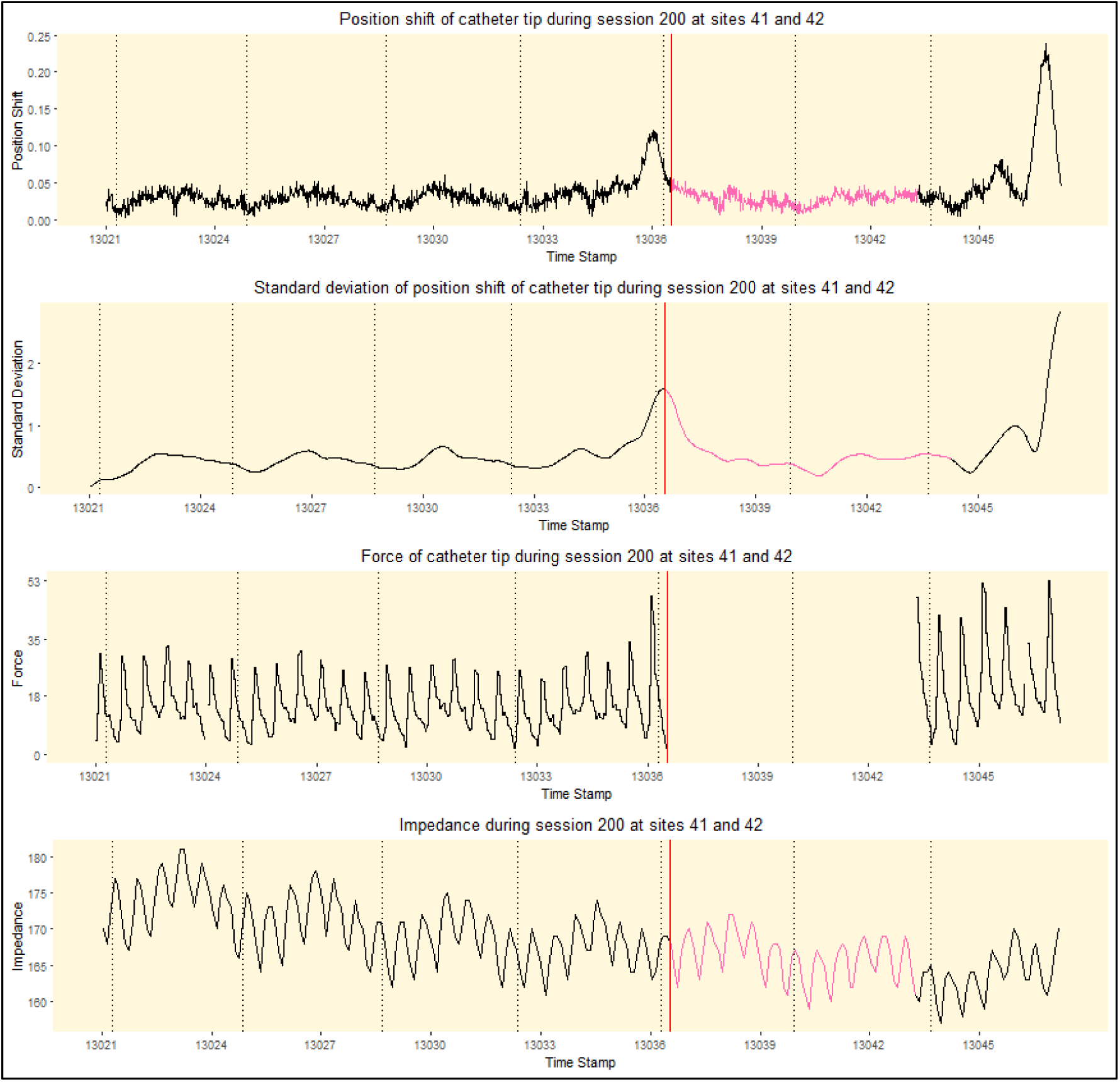
Reconstituted curves from VISITAG™ Module data export demonstrating first-annotated site “end” time-point (i.e. site 1-to-2 transition) at the first 0g CF event, 15.5s (ACCURESP™ “on”) and 15.6s (ACCURESP™ “off”) following RF onset in case 5, right PV. This small difference in the annotated RF duration between ACCURESP™ settings results in a single vertical red line representing ACURESP™ “on” and “off” annotation timing. Catheter tip position shift, (position) standard deviation (SD), CF and impedance are plotted separately; x-axis displays the PIU time stamp (i.e. running case time, shown in thousands of seconds) and vertical dashed lines indicate end-expiration time-points. The force-over-time 100% minimum 1g CF filter ensures that automated RF annotation only occurs when CF is continuously maintained ≥1g, indicated by black curves; pink position shift, SD and impedance curves indicate <100% CF filter preference attainment. The SD curve returns to black at 1s (or 60 sites) following attainment of both CF and position stability within the chosen VISITAG™ Module filter preferences, in view of the CARTO^®^3 SD calculation “system logic”. Note: “Session 200 at sites 41 and 42” represents the unique automatically generated VISITAG™ Module annotated identifiers for this site.

**Figure 2:**
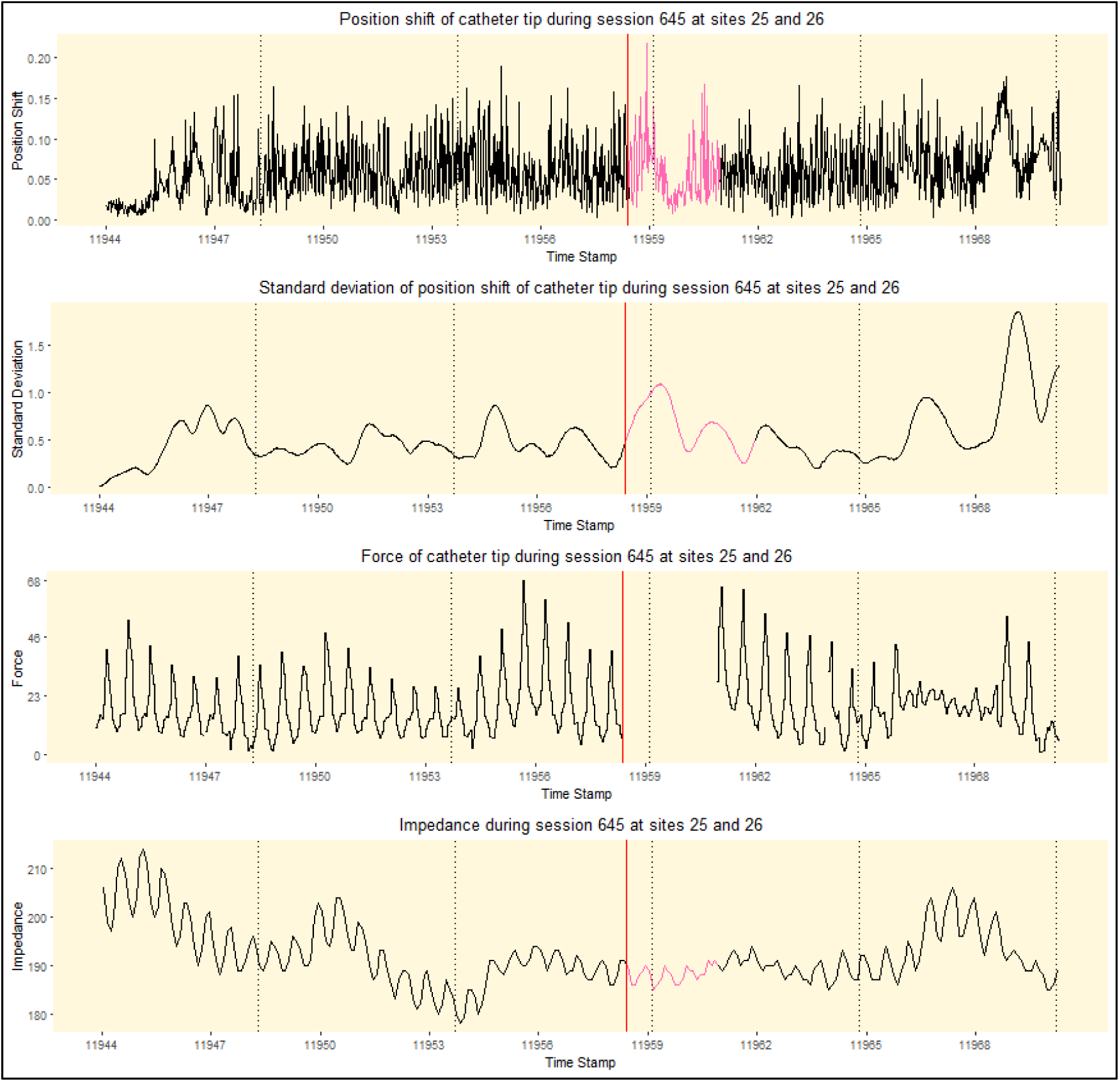
Reconstituted curves from VISITAG™ Module data export demonstrating first-annotated site “end” time-point (i.e. site 1-to-2 transition) at the first 0g CF event, 14.4s following RF onset in case 10, right PV. This small difference in the annotated RF duration between ACCURESP™ settings results in a single vertical red line representing ACURESP™ “on” and “off” annotation timing; other plot elements are as per figure 1. Note: “Session 645 at sites 25 and 26” represents the unique automatically generated VISITAG™ Module annotated identifiers for this site.

**Figure 3:**
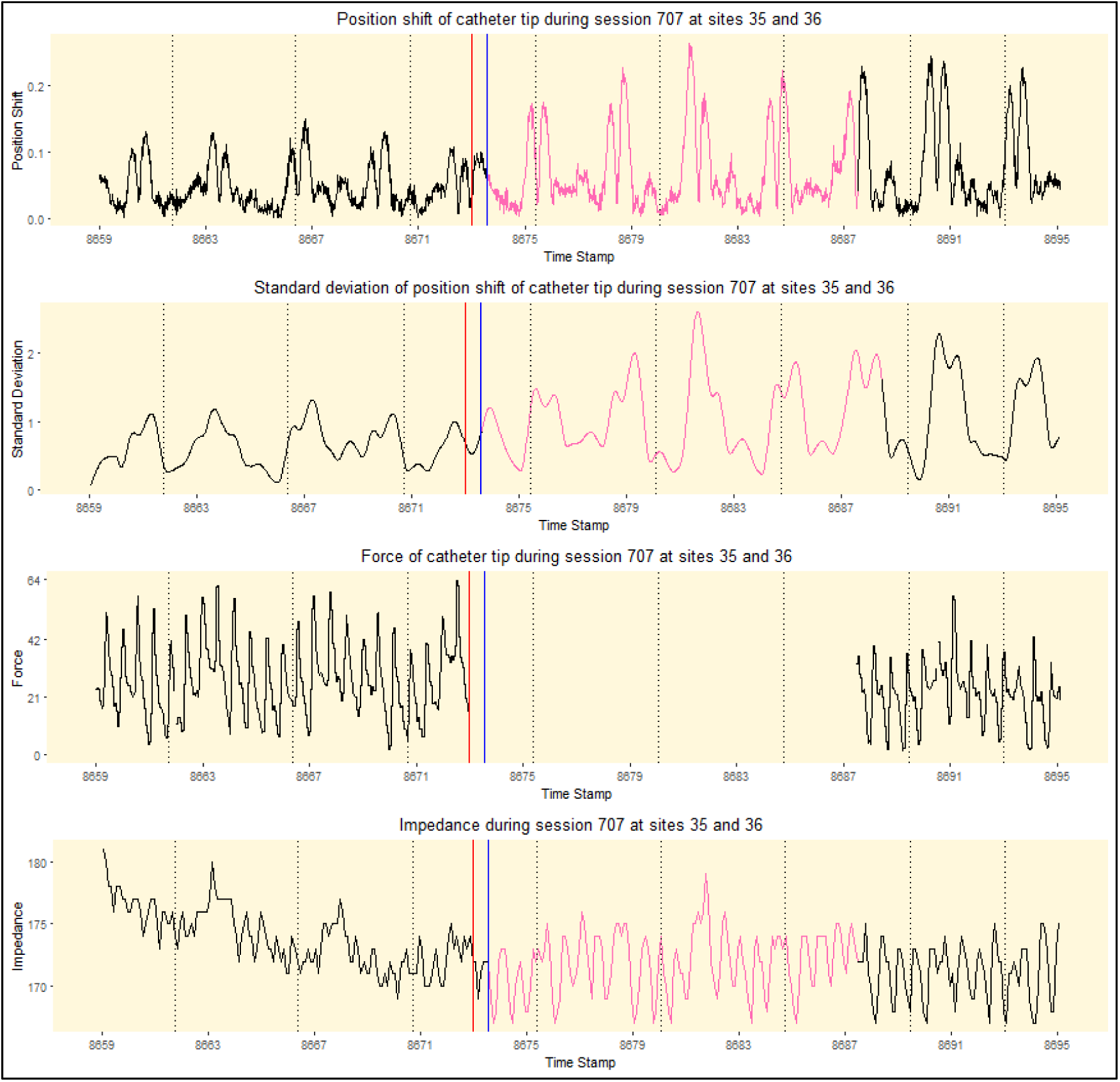
Reconstituted curves from VISITAG™ Module data export demonstrating first-annotated site “end” time-point (i.e. site 1-to-2 transition) at the first 0g CF event, 14.0s (ACCURESP™ “on”, red line) and 14.6s (ACCURESP™ “off”, blue line) following RF onset in case 11, right PV; other plot elements are as per figure 1. Note: “Session 707 at sites 35 and 36” represents the unique automatically generated VISITAG™ Module annotated identifiers for this site.

Four site 1-to-2 annotated transitions were effected with constant catheter-tissue contact and associated with UE morphology change from pure R (site 1 completion) to RS (site 2 onset). In this group, the maximum difference (ACCURESP™ “on” minus “off”) in RF duration, FTI, ILD and impedance drop was 11.3s, 139g.s, 2.6mm and 3.3Ω respectively (table 3), with the first indication of catheter movement represented by ACCURESP™ “off” annotation in all cases (figures 4-7). The greatest difference was seen in a case where site 1-to-2 ILD with ACCURESP™ “off” was 4.1mm (reconstituted data curves, figure 4); at 15.2s following RF onset there was an abrupt increase in catheter position shift and SD, with a corresponding change in CF waveform, yet while the blue vertical line indicating annotation site 1-to-2 transition according to ACCURESP™ “off” coincided with these changes and the “per protocol” ∼15s RF at site 1, the red vertical line indicating annotated site 1-to-2 transition using ACCURESP™ “on” is shown occurring 11.3s later.

**Figure 4:**
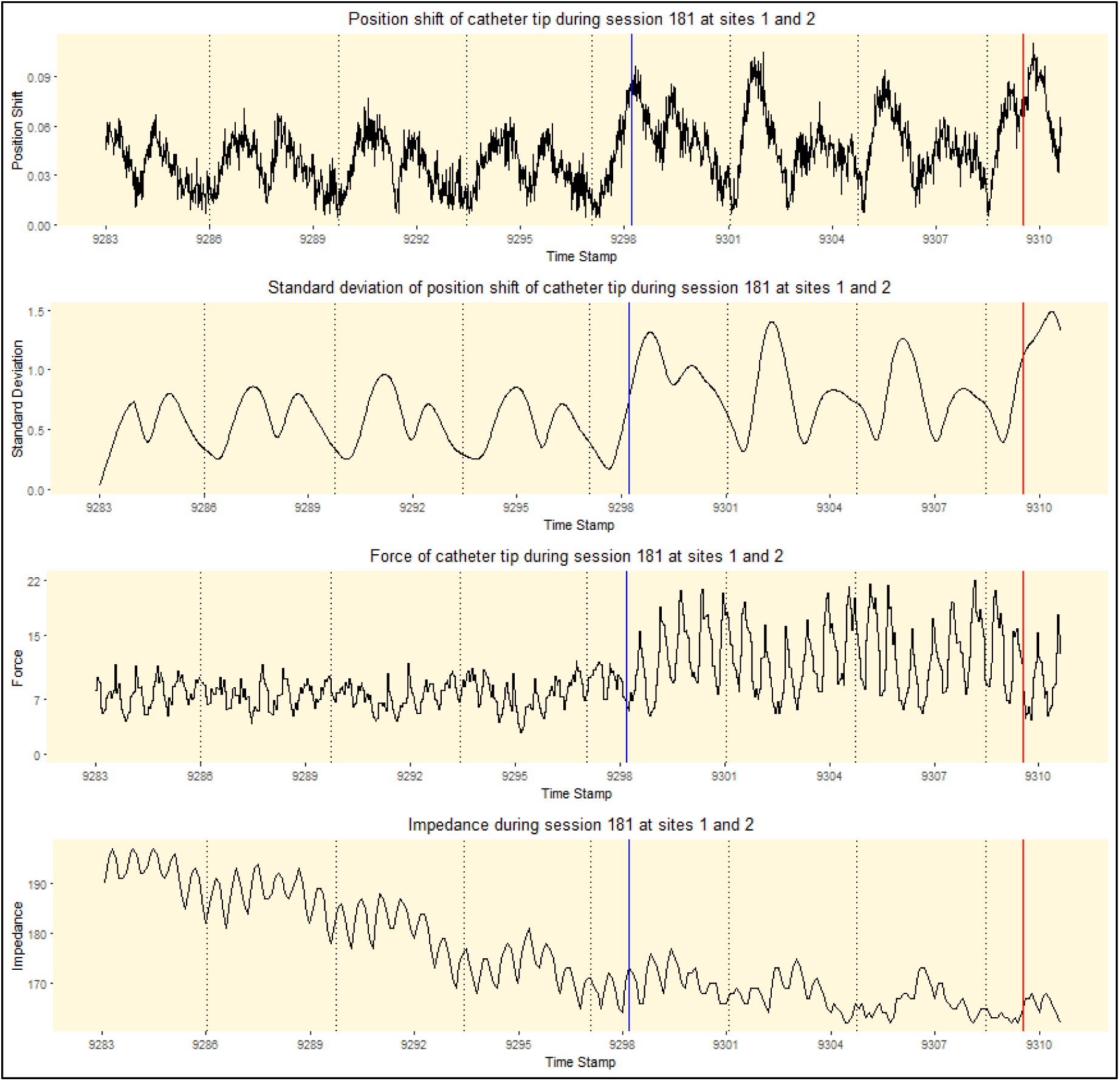
Reconstituted curves from VISITAG™ Module data export demonstrating first-annotated site “end” time-point (i.e. site 1-to-2 transition) at 15.2s (ACCURESP™ “off”, blue line) and 26.5s (ACCURESP™ “on”, red line) following RF onset in case 5, left PV. Transition to site 2 with ACCURESP “off” coincided with UE morphology change from pure R at site 1 completion, to RS at site 2 onset (ACCURESP™ “off” ILD 4.1mm); clear changes in the catheter tip position shift, (position) standard deviation (SD) and CF coincide with ACCURESP™ “off” annotation. All curves are drawn black, since CF was maintained ≥1g and the catheter movement was sufficiently rapid to ensure that all catheter tip location data was annotated to either site 1, or 2 (ACCURESP™ “off”). Note: “Session 181 at sites 1 and 2” represents the unique automatically generated VISITAG™ Module annotated identifiers for this site.

**Figure 5:**
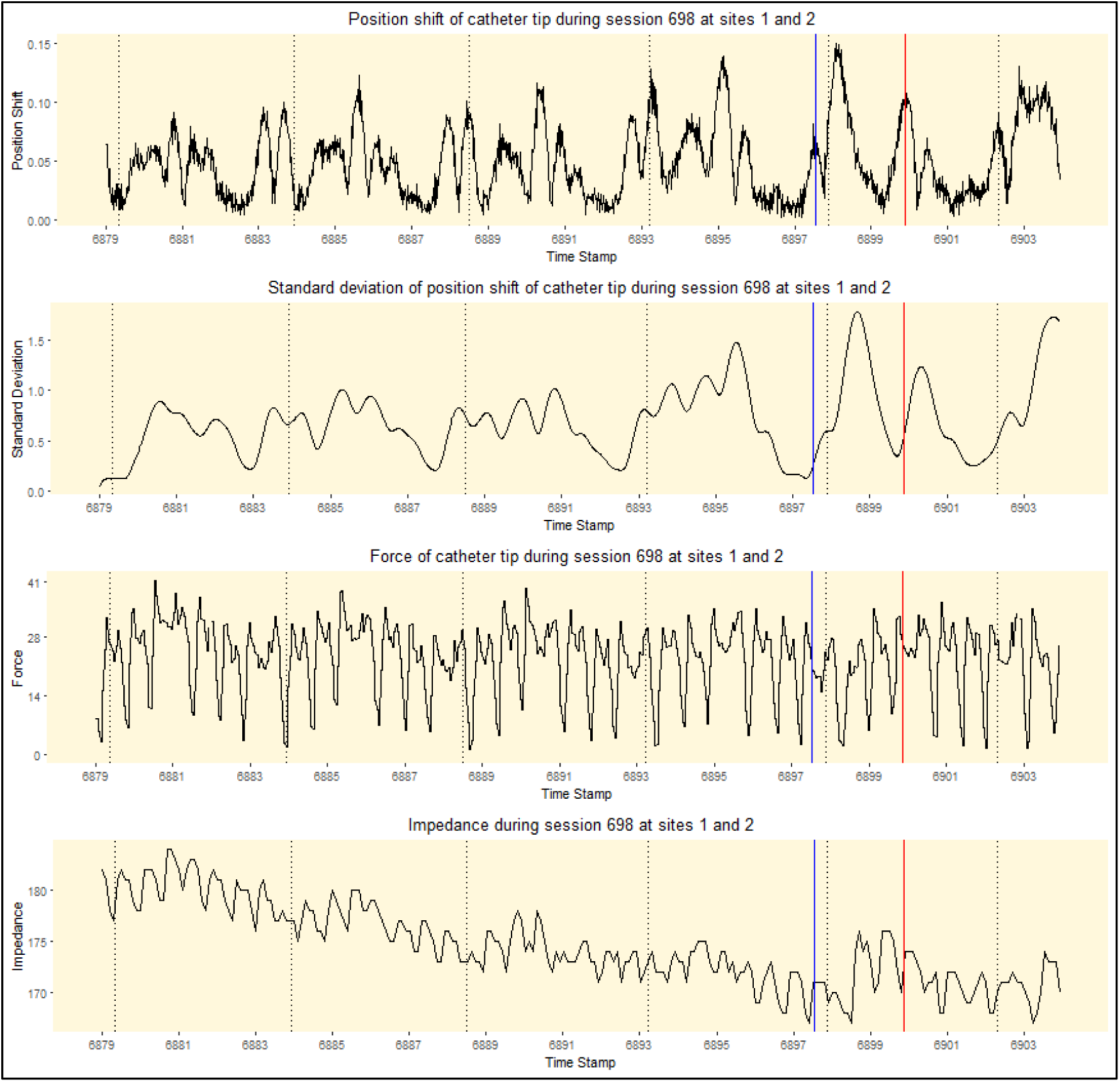
Reconstituted curves from VISITAG™ Module data export demonstrating first-annotated site “end” time-point (i.e. site 1-to-2 transition) at 18.6s (ACCURESP™ “off”, blue line) and 20.9s (ACCURESP™ “on”, red line) following RF onset in case 11, left PV. Transition to site 2 with ACCURESP “off” coincided with UE morphology change from pure R at site 1 completion, to RS at site 2 onset (ACCURESP™ “off” ILD 5.2mm). The greatest position shift value is seen occurring closest to ACCURESP™ “off” annotation timing; other plot elements are as per figure 4. Note: “Session 698 at sites 1 and 2” represents the unique automatically generated VISITAG™ Module annotated identifiers for this site.

**Figure 6:**
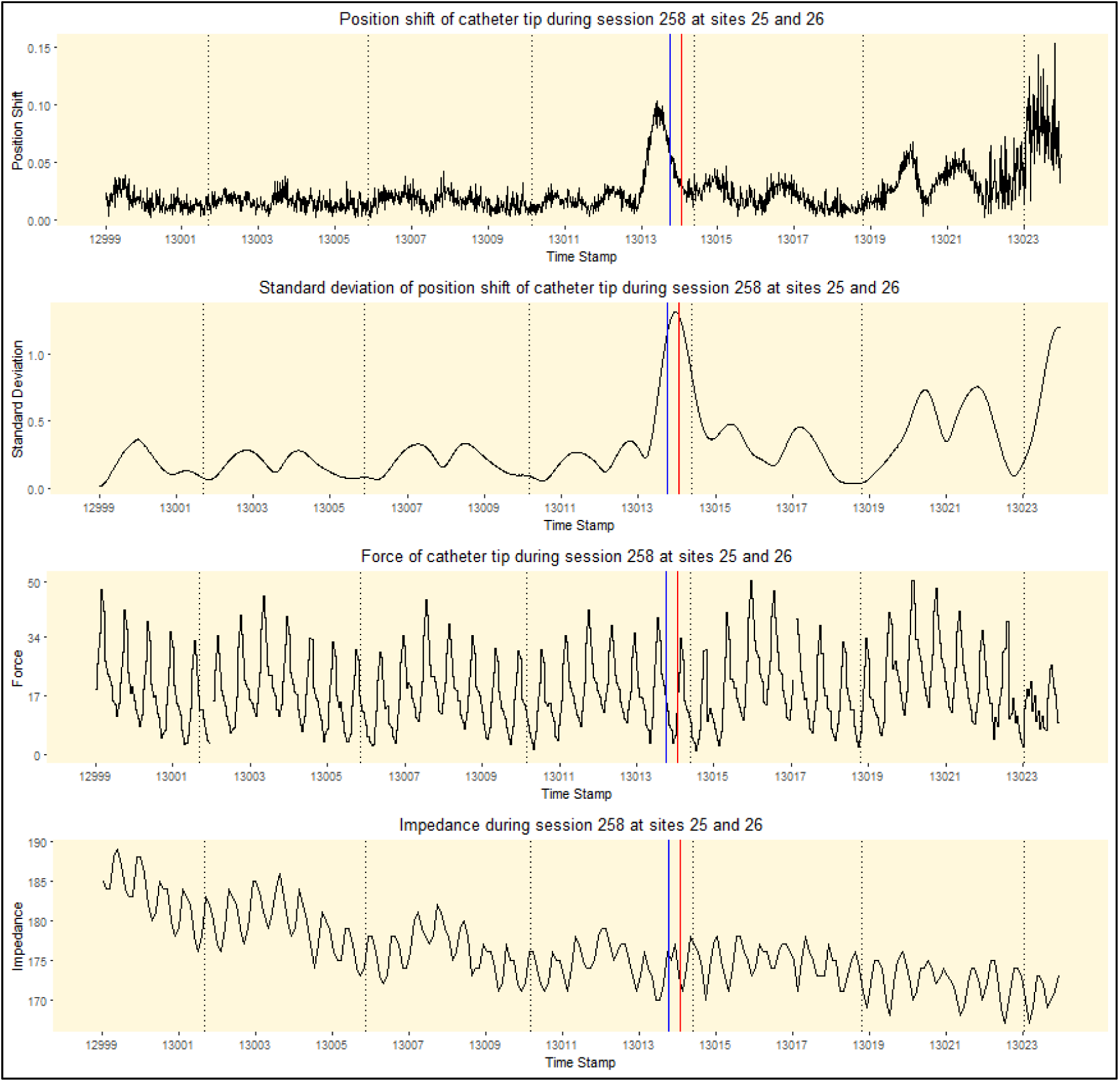
Reconstituted curves from VISITAG™ Module data export demonstrating first-annotated site “end” time-point (i.e. site 1-to-2 transition) at 14.8s (ACCURESP™ “off”, blue line) and 15.1s (ACCURESP™ “on”, red line) following RF onset in case 22, right PV. Transition to site 2 with ACCURESP “off” coincided with UE morphology change from pure R at site 1 completion, to RS at site 2 annotation onset (ACCURESP™ “off” ILD 5.0mm); clear changes in the catheter tip position shift and (position) standard deviation (SD) occur just before ACCURESP™ “off” annotation. All curves are drawn black, since CF was maintained ≥1g and the catheter movement was sufficiently rapid to ensure that all catheter tip location data was annotated to either site 1, or 2 (ACCURESP™ “off”). Note: “Session 258 at sites 25 and 26” represents the unique automatically generated VISITAG™ Module annotated identifiers for this site.

**Figure 7:**
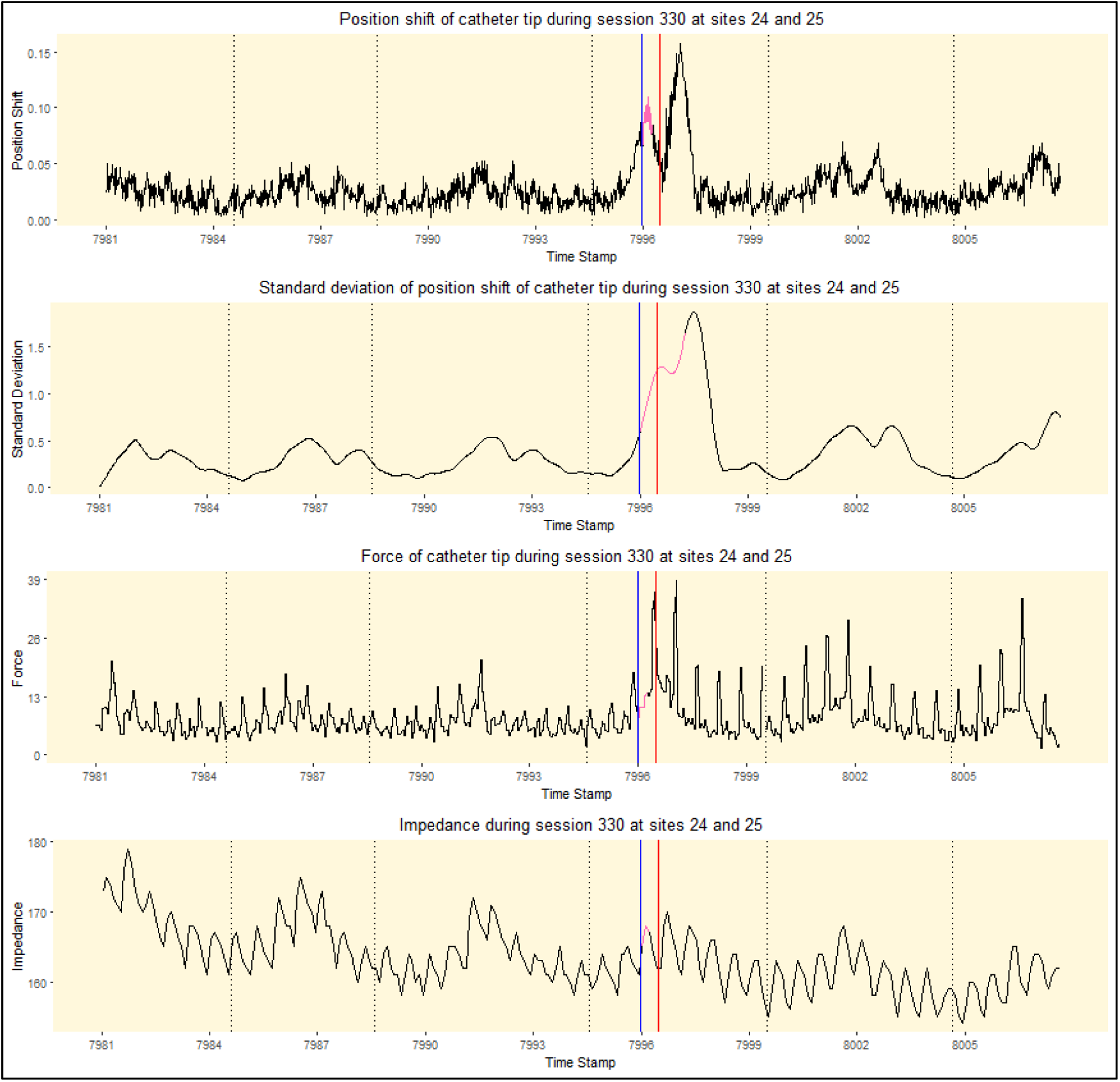
Reconstituted curves from VISITAG™ Module data export demonstrating first-annotated site “end” time-point (i.e. site 1-to-2 transition) at 15.0s (ACCURESP™ “off”, blue line) and 15.5s (ACCURESP™ “on”, red line) following RF onset in case 23, right PV. Transition to site 2 with ACCURESP “off” coincided with UE morphology change from pure R at site 1 completion, to RS at site 2 annotation onset (ACCURESP™ “off” ILD 8.8mm); the first obvious changes in the catheter tip position shift, (position) standard deviation (SD) and CF coincide with ACCURESP™ “off” annotation. The curves are briefly shown in pink since with ACCURESP™ “off” there was 0.28s of non-annotated inter-ablation site transition time due position instability >2*(2mm SD) according to the CARTO^®^3 “system logic”; the position SD curve is pink for a total of 1.28s, reflecting system SD calculations over a minimum of 60 positions (i.e. 1s). Note: “Session 330 at sites 24 and 25” represents the unique automatically generated VISITAG™ Module annotated identifiers for this site.

The remaining nine site 1-to-2 transitions with CF ≥1g were associated with continuous pure R UE morphology – i.e. at both site 1 end and site 2 onset. Comparison of annotated biophysical data (ACCURESP™ “on” minus “off”) demonstrated a difference in timing of site 1 RF duration of >1s in eight; range −1.3 – 8.6s, mean 3.7 [SD: 4.0] s (table 1, data supplement). In this group, the maximum difference in site 1 annotated RF duration, FTI, ILD and impedance drop was 8.6s, 208g.s, 7.7mm and 1.4Ω respectively; ACCURESP™ “on” resulted in greater values for annotated data in seven of nine site 1-to-2 transitions. However, considering multiple measures of catheter position stability, the first indication of catheter motion was appropriately identified with ACCURESP™ “off” in eight transitions; only one demonstrated a difference in annotation timing without clinical importance (i.e. 0.1s, supplementary figure 8).

**Figure 8:**
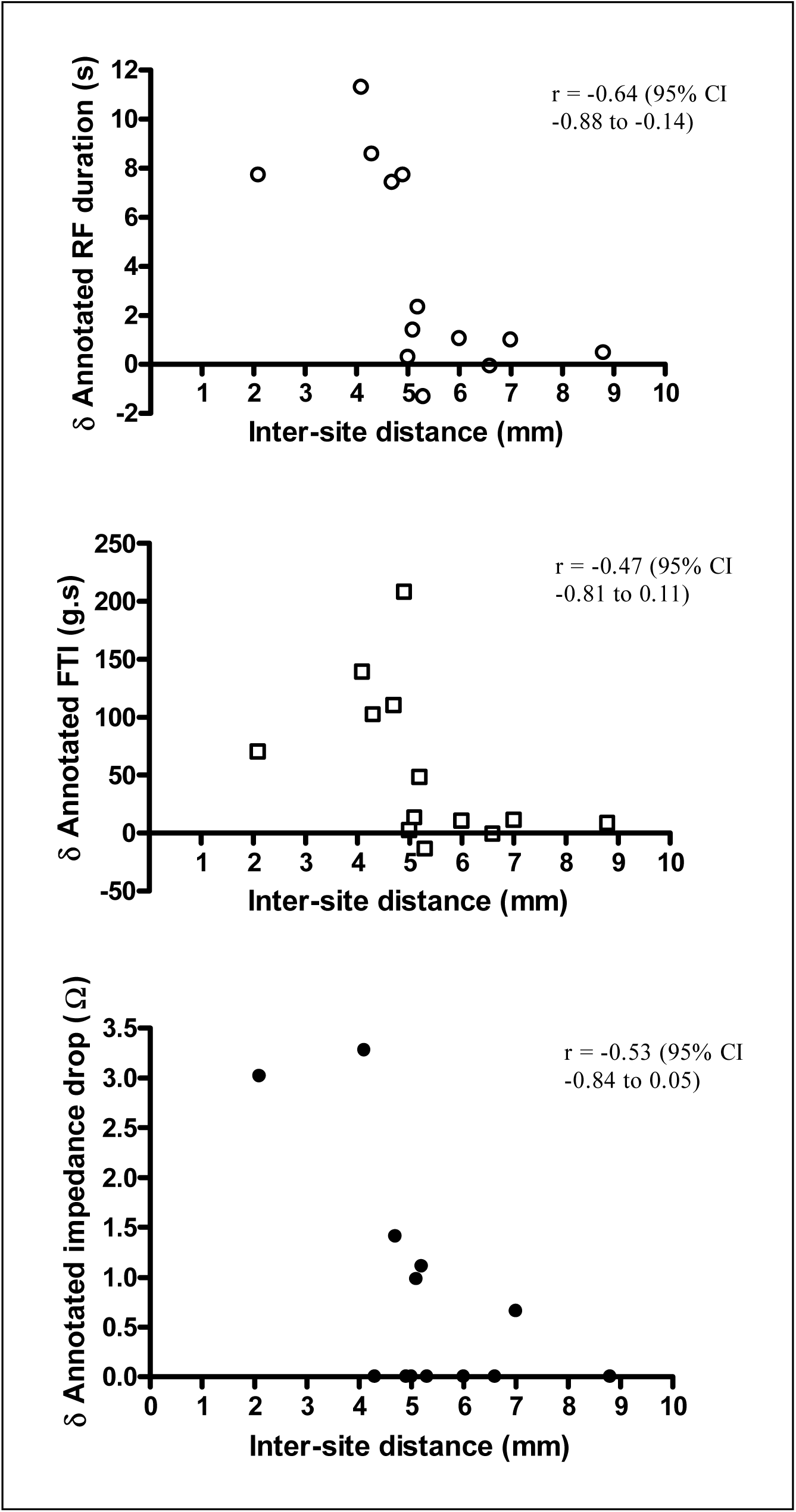
ACCURESP™ “on” minus “off” difference in annotated RF duration, FTI and impedance drop according to site 1-2 inter-ablation site distance with ACCURESP™ “off”. Data are shown for all 13 LAPW inter-ablation site transitions achieved during constant catheter-tissue contact, with corresponding correlation data.

For all thirteen site 1-to-2 transitions achieved with constant catheter-tissue contact, the relationship between differences in annotated RF data (ACCURESP™ “on” minus “off”) and ILD (with ACCURESP™ “off”) is shown in figure 8. There was a strong negative correlation between the difference in annotated RF duration and ILD – Pearson r −0.68 (95% confidence interval (CI) −0.91 to −0.13, p=0.02, figure 8). Consequently, while the maximum difference in annotated RF duration with site 1-to-2 ILD ≥6mm was 1.1s, an ILD ≤5mm was associated with maximal difference in annotated RF duration of 11.3s. There was a moderate negative correlation between the difference in annotated FTI and ILD – Pearson r −0.47 (95% CI −0.81 to 0.11, p=0.10) and a moderate negative correlation between the difference in impedance drop and ILD – Pearson r −0.53 (95% CI −0.84 to 0.05, p=0.07).

Data supplement figures 2 – 10 demonstrate all remaining annotated site 1-to-2 transitions, with corresponding position shift, SD, CF and impedance data.

### Analyses at subsequently annotated LAPW sites during continuous RF delivery

All left-sided LAPW lesion sets and 6 of 8 right-sided LAPW lesion sets were completed during continuous, uninterrupted RF application; the onset timing of transitions for sites 2-to-3, 3-to-4 and 4-to-5 is shown in table 4. For left-sided LAPW lesions there was a progressively greater difference in timing between annotated sites of transition with ACCURESP “on” versus “off”, increasing from a median of 1.2s (IQR 0.4 – 5.0) at site 2 onset, to a median of 14.6s (IQR 7.6 – 23.0) at site 5 annotation onset. Examples for one case are shown in supplementary figures 10-13 with annotated position shift, SD, CF and impedance data. For right-sided LAPW lesions annotated during continuous RF application there was no progressive increase in the difference between ACCURESP “on” versus “off” annotation. However, for right-sided sites, 35% (95% CI 21-53) annotated transitions occurred with 0g CF, whereas for left-sided sites this was only 3% (95% CI 0 – 17).

**Table 4:** Difference in onset timing (ACCURESP™ “on” minus “off”) of consecutively annotated LAPW sites during continuous RF according to the PIU “system time”; median (1^st^ – 3^rd^ quartile, IQR), maximum and minimum differences are shown.

| Annotated site<br>number | Left-sided annotation onset difference (s) |  |  | Right-sided annotation onset difference (s) |  |  |
| --- | --- | --- | --- | --- | --- | --- |
|  | (N=8) |  |  | (N=6) |  |  |
|  | *Median (IQR)* | *Maximum* | *Minimum* | *Median (IQR)* | *Maximum* | *Minimum* |
| 2 | 1.2<br>(0.4 – 5.0) | 11.3 | -1.3 | 0.3<br>(-0.02 – 8.0) | 8.6 | -0.02 |
| 3 | 7.3<br>(4.4 – 10.8) | 16.3 | 0.7 | 4.2<br>(-0.04 – 10.8) | 11.9 | -0.04 |
| 4 | 11.6<br>(6.3 – 14.4) | 19.5 | 0.02 | -0.04<br>(-0.9 – 6.9) | 13.4 | -1.7 |
| 5 | 14.6<br>(7.6 – 23.0) | 28.2 | 2.1 | 2.5<br>(0.8 – 11.9) | 15.6 | -0.06 |

## Discussion

The main findings of this present study are as follows: (1) ACCURESP™ respiratory adjustment resulted in important differences in VISITAG™ Module annotation at the LAPW, with ACCURESP™ set “on” overall resulting in significantly greater per-site RF duration and ILD; (2) At sites of deliberate catheter motion between first and second-annotated sites effected without loss of catheter-tissue contact, annotated transition using ACCURESP™ “off” demonstrated suitable accuracy towards catheter motion detection and inter-ablation site transition annotation in all cases. However, ACCURESP™ “on” resulted in inappropriate annotation delay of ≥1s in the majority (i.e. 9 of 13 site 1-to-2 annotated transitions); (3) ACCURESP™ setting was only without significant effect on RF annotation timing when sites of deliberate catheter motion were accompanied by loss of catheter-tissue contact (i.e. 0g CF) and while using a force-over-time filter of 100% minimum 1g; (4) There was a significant inverse relationship between the difference in first-annotated site RF duration (ACCURESP™ “on” minus “off”), and the ACCURESP™ “off” site 1-to-2 ILD. Consequently, with ACCURESP™ “on”, catheter tip motion of up to 7mm was not immediately identified, resulting in delayed annotation of site 1-to-2 transition and important error in per-site RF parameters. More simply, ACCURESP “on” may effectively render an operator “blind” to the immediate occurrence of small but clinically important catheter displacement events.

These results may be better understood when considering the VISITAG™ Module annotation “system logic” and how this is modified by ACCURESP™ use. Briefly, the ablation catheter tip position is measured in a “rolling window” of 60 sites per second (i.e. intervals of 16/17ms), from which is calculated the standard deviation (SD). With ACCURESP™ “off”, RF annotation occurs when both every second of position data is within twice (2x) the user-defined position SD, and a total consecutive minimum of 3s is within (1x) the SD. However, with ACCURESP™ “on”, the position stability filter operates over a minimum of two respiratory cycles, using position data “adjusted” to end-expiration; here, automated ablation site annotation occurs when these position stability targets are met *following data adjustment*. Importantly, annotation only occurs when CF filter preferences are also satisfied. However, in contrast to position stability filtering, the CF filter is applied continuously regardless of ACCURESP™ setting (i.e. independent of the respiratory cycle). Therefore, with the force-over-time 100% minimum 1g CF filter used during this present report, RF annotation “end” logic is fulfilled by any 0g CF event at any stage in the respiratory cycle, with ACCURESP™ both “on” and “off”.

### Importance of these findings: Historical perspective

Following the advent of VISITAG™ Module automated RF annotation, this tool has been demonstrated to facilitate both the derivation of hypothetically ideal per-site ablation parameters and their subsequent delivery during CF and VISITAG™ Module-guided PVI.^6,7,13,15^ Such targets can only be considered appropriate when “derivation phase” methodology employed a suitable definition for a stable site of catheter-tissue interaction during RF application. Notwithstanding the theoretical difficulty resulting from a choice of CF filter permitting intermittent catheter-tissue contact (i.e. by definition a stable site can only occur in the setting of constant tissue contact), the foundational study supporting a regional difference in ablation target values also used ACCURESP™ “on” in all cases; procedures performed under GA and with IPPV failing to trigger ACCURESP™ had the tidal volume increased to ensure ACCURESP™ triggering and “on” setting use (Molloy Das, personal communication).^13^ As this present report has demonstrated that catheter tip movements of up to 7mm are not immediately identified with ACCURESP™ “on”, these previously identified ablation targets are likely to be importantly flawed.

### Clinical importance in light of novel high power, short per-site RF duration protocols

RF lesion formation occurs more rapidly at higher power.^17^ Recently reported high power short duration (HPSD) protocols (50W, typically ∼5s per site)^18,19^ are particularly attractive for operators wishing to move away from the requirement to maintain a stable catheter position for the ∼20-40s previously considered necessary to achieve TM RF effect during PVI.^1,20^ A greater rate of lesion formation means HPSD protocols are particularly dependent upon suitable methodology towards the immediate and accurate determination of clinically relevant catheter position instability. However, to our knowledge no HPSD manuscript includes details of whether respiratory adjustment was applied to RF annotation logic. Alongside the recommendation (from Biosense Webster) for routine VISITAG™ Module use with ACCURESP™ “on”, this calls into question the validity of such HPSD “per-site” ablation targets (since they were likely to have been performed using RF annotation with respiratory adjustment) and whether present HPSD protocols may be considered reproducible.

### Future directions for research

For VISITAG™ Module-guided procedures, the other important determinant of RF annotation logic towards a suitable definition of a stable site of catheter tissue interaction during RF application is the choice of position stability filter setting. Indeed, a possible solution to the inaccuracy of ACCURESP™ “on” as described in this present report is to use a smaller position stability filter range – e.g. 1.5mm. Although a complete description of the effects of this position filter setting is beyond the scope of this present report, data supplement figure 15 demonstrates annotated left PV site 1 data for case #5 (reconstituted data curves in figure 4), using the VISITAG™ Module preference settings ACCURESP™ “off” with 2mm position stability, and ACCURESP™ “on” with 1.5mm position stability range. Importantly, when using 1.5mm range with ACCURESP™ “on”, annotated RF duration at site 1 is 1.6s greater, yet as shown in figure 4, position stability and CF changes indicating first catheter displacement coincide with annotation using ACCURESP™ “off” and 2mm range (blue vertical line). In this case, further evidence towards this earlier time point representing the “true” site of catheter displacement is provided by the UE morphology change from pure R (site 1 end) to RS (site 2 onset) coinciding with ACCURESP™ “off” and 2mm range annotation timing. Such a delay of 1.6s is likely to be clinically important, particularly when considering HPSD ablation.

Together with a steerable sheath for catheter support, high frequency jet ventilation (HFJV) has been shown to reduce the occurrence of acute and chronic pulmonary vein reconnections as well as improve freedom from AF.^21^ Theoretically, complete absence of respiratory motion not only eliminates the requirement for respiratory adjustment, but also confers the advantage of eliminating a requirement for adjustments to sheath and/or catheter position according to the respiratory cycle; presently, there is no means to capture data on such patient and/or operator-specific movements. Accordingly, VISITAG™ Module and CF-guided PVI protocols using HFJV may demonstrate greater reproducibility and efficacy.

### Limitations

This report was of a single operator’s practice, with analyses limited to LAPW lesions in view of the risk of atrio-oesophageal fistula resulting from excessive RF application at this site. It is possible that ACCURESP™ settings demonstrate greater annotation concordance at alternative left atrial sites; such analyses were beyond the scope of this present report. The personalised VISITAG™ Module filter preferences (force-over-time 100% minimum 1g, with 2mm position stability), overdrive atrial pacing and steerable sheath use towards achieving catheter stability and optimal ILD during continuous RF application must be taken into account when considering these experimental findings. If the more commonly employed force-over-time 30% minimum 4-5g CF filter was employed^7,13^, intermittent catheter-tissue contact would only trigger per-site annotation “end” if position data breached the chosen (position) stability criteria at that time-point. We elected not to perform a complete re-analysis of exported data with this different CF filter setting, since a site of stable catheter-tissue interaction during RF by definition may only occur in the setting of constant catheter-tissue contact (i.e. assuming no catheter CF measurement error, force-over-time 100% ≥1g). Also, a complete description of the effects of changing the position stability filter setting is beyond the scope of this present report; this will be the subject of a future manuscript.

Catheter tip motion characteristics following deliberate movement may importantly differ from unintentional events; due to the very high attainment of target per-site RF delivery in this present report, a description of position stability, CF and impedance profiles at sites of accidental catheter motion is beyond the scope of this present report. However, when deliberate catheter displacement events of up to ∼5-7mm were not immediately identified when using annotation with ACCURESP™ “on”, similar degrees of movement during unintentional catheter displacement events are likely to be missed.

The findings of this present report can only be directly applied to VISITAG™ Module-guided PVI. However, EnSite Precision™ (Abbott) – the only other system with an automated RF annotation module, AutoMark™ – also utilises respiratory motion compensation logic. Importantly, EnSite Precision™ employs permanent RF data and electrogram storage; accordingly, the findings of this present report may help stimulate retrospective investigations into the methodological rigour of different AutoMark™ annotation settings during AF ablation.

It remains impossible to determine the site-specific magnitude of any out-of-phase catheter tissue interaction occurring during constant contact using the methodology described in this present report; intra-cardiac echo was never used. Therefore, even with ACCURESP™ “off”, respiratory motion may represent an important determinant of the recently identified heterogeneity in RF effect during PVI.^9,22,23^ Furthermore, cardiac cycle-induced motion may represent another important determinant of catheter instability, although atrial overdrive pacing during RF delivery has been demonstrated to improve catheter stability and impedance reduction;^24^ this technique was also used in this present report.

Finally, this report is based on analyses of 8 PVI procedures from a total cohort of 25 patients – i.e. a small sample size from which to conventionally derive meaningful data. However, RF annotation-guided ablation represents a very “data-rich” operative environment and analyses for each ∼15s first-site RF application alone utilise 3090 exported data points (i.e. CF at 20Hz, impedance at 10Hz, and both position SD and position shift at 60Hz). Therefore, although drawn from an eight-patient cohort, the findings of this present report represent a total analysis of ∼165,000 data points.

## Conclusions

During CF and VISITAG™ Module annotation-guided PVI, ACCURESP™ respiratory adjustment results in important delays to the identification and annotation of sites of deliberate catheter motion of up to 5-7mm at the LAPW. Therefore, previously derived ablation targets using ACCURESP™ set “on” may be importantly flawed, and on-going respiratory adjustment use is likely to represent an important impediment towards greater PVI procedural reproducibility, efficacy and safety. In contrast, RF annotation with ACCURESP™ set “off” demonstrated suitable catheter motion detection capabilities. Based on these findings, we recommend setting ACCURESP™ “off” during VISITAG™ Module and CF-guided PVI.

## Acknowledgements

I am grateful to Cherith Wood, Daniel Newcomb and Ian Lines, Cardiac Physiologists, for their technical support into all cases conducted during this report. I am also grateful to Robert Pearce and Vicky Healey (Biosense Webster Inc.) for additional technical assistance and to Noam Seker-Gafni, Tal Bar-on, Einav Geffen, Assaf Rubissa and colleagues at the Haifa Technology Center, Israel (Biosense Webster Inc.) for their help with VISITAG™ Module technical queries. I would also like to thank Ieuan Coombes, Gary Smith, Kristi Tanouye, Travis Dahlen and Mark Hagfors (Abbott Inc.) for clarifying EnSite Precision™ and AutoMark™ system elements.

## Sources of Funding

I am grateful to the “Sarkar Research and Training” charitable fund, University Hospitals Plymouth NHS Trust for a donation of £1000, funding the R software code development and extended data analyses by teams at the Department of Medical Statistics, Plymouth University Peninsula Schools of Medicine and Dentistry.

## Disclosures

None

