## Supplementary data for "Ablation catheter motion detection during contact force and VISITAG™ Module-guided pulmonary vein isolation"

**Detection of catheter displacement during contact force and VISITAG™ Module-guided pulmonary vein isolation: An improved approach towards respiratory adjustment**

Authors:	David R. Tomlinson BM BSc MD^1^, Katie Biscombe^2^, John True BSc MSc^2^, Joanne Hosking BSc PhD^2^ and Adam J. Streeter MSc PhD^2^

Corresponding address:	^1^University Hospitals Plymouth NHS Trust, South West Cardiothoracic Centre, Derriford Hospital, Plymouth, UK, PL6 8DH

^2^Department of Medical Statistics, Plymouth University Peninsula Schools of Medicine and Dentistry, ITTC Building 1, Plymouth Science Park, Plymouth, Devon PL6 8BX, UK.

| Case number | PV | Site 1 RF duration (s) | | | Distance to site 2 (mm) | | | Site 1 impedance drop (Ω) | | | Site 1 FTI (g.s) | | |
| --- | --- | --- | --- | --- | --- | --- | --- | --- | --- | --- | --- | --- | --- |
|  |  | ACCURESP | | | ACCURESP | | | ACCURESP | | | ACCURESP | | |
|  |  | ON | OFF | *Diff* | ON | OFF | *Diff* | ON | OFF | *Diff* | ON | OFF | *Diff* |
| 2 | Left | 16.5 | 15.5 | *1.0* | 7.7 | 6.0 | *1.7* | 13.4 | 13.4 | *0* | 204 | 194 | *10* |
| 2 | Right | 23.7 | 16.3 | *7.4* | 6.3 | 4.7 | *1.6* | 8.2 | 6.8 | *1.4* | 251 | 141 | *110* |
| 10 | Left | 14.4 | 15.7 | *-1.3* | 7.0 | 5.3 | *1.7* | 10.8 | 10.8 | *0* | 156 | 170 | *-14* |
| 14 | Left | 15.9 | 14.5 | *1.4* | 6.3 | 5.1 | *1.2* | 17.8 | 16.8 | *1.0* | 169 | 156 | *13* |
| 14 | Right | 23.6 | 15.9 | *7.7* | 12.6 | 4.9 | *7.7* | 8.8 | 8.8 | *0* | 590 | 382 | *208* |
| 16 | Left | 15.6 | 7.9 | *7.7* | 6.5 | 2.1 | *4.4* | 8.5 | 5.5 | *3* | 144 | 74 | *70* |
| 16 | Right | 23.8 | 15.2 | *8.6* | 6.6 | 4.3 | *2.3* | 12.9 | 12.9 | *0* | 369 | 267 | *102* |
| 22 | Left | 14.8 | 14.9 | *-0.1* | 7.0 | 6.6 | *0.4* | 13.6 | 13.6 | *0* | 200 | 201 | *-1* |
| 23 | Left | 16.0 | 15.0 | *1.0* | 6.2 | 7.0 | *-0.8* | 42.6 | 42.0 | *0.6* | 139 | 128 | *11* |

Supplementary table 1: Annotated biophysical data at sites of catheter position instability occurring at the transition from first to second-annotated LAPW sites with pure R UE morphology present at both site 1 completion and site 2 onset. Annotated RF duration, site 1-2 ILD, impedance drop and force time integral (FTI) data are displayed according to ACCURESP™ setting, with the difference (“Diff” – i.e. ACCURESP™ setting “on” minus “off”) also shown (PV, pulmonary vein).

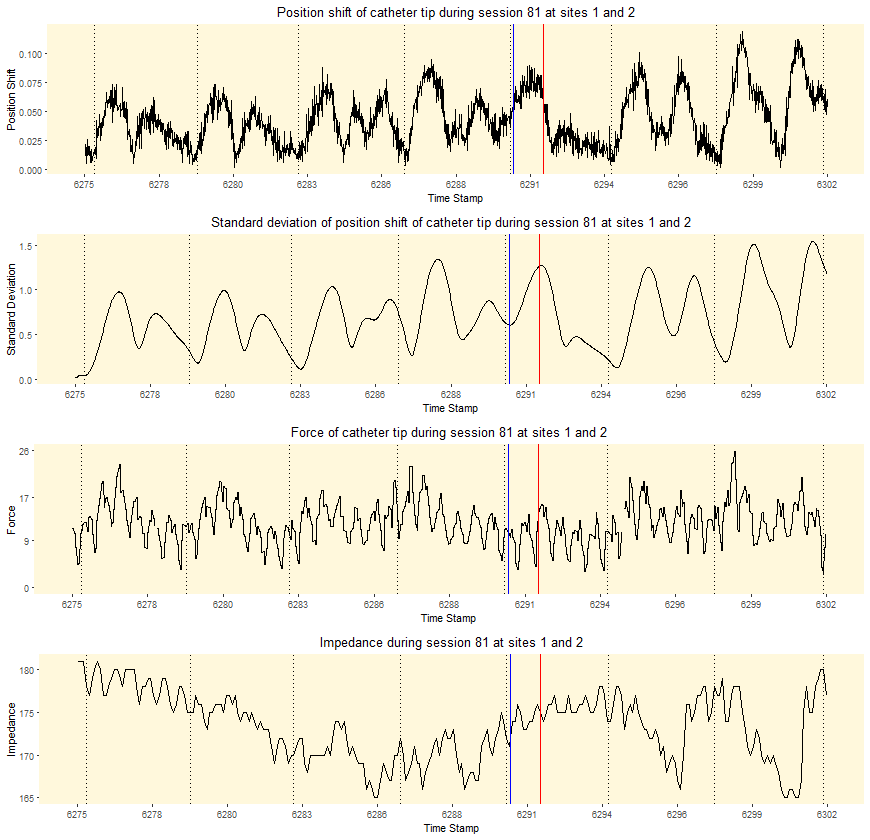

Supplementary figure 1: Reconstituted curves from VISITAG™ Module data export demonstrating first-annotated site “end” time-point at 15.5s (ACCURESP™ “off”, blue line) and 16.5s (ACCURESP™ “on”, red line) following RF onset in case 2, left PV. Inter-ablation site transition occurred with pure R UE morphology present at both site 1 completion and site 2 onset (ACCURESP™ “off” ILD 6.0mm); subtle changes in the CF and impedance curves occur concurrent with annotated site transition. All curves are drawn black, since CF was maintained ≥1g and the catheter movement was sufficiently rapid to ensure that all catheter tip location data was annotated to either site 1, or 2 (ACCURESP™ “off”). Note: “Session 81 at sites 1 and 2” represents the unique automatically generated VISITAG™ Module annotated identifiers for this site.

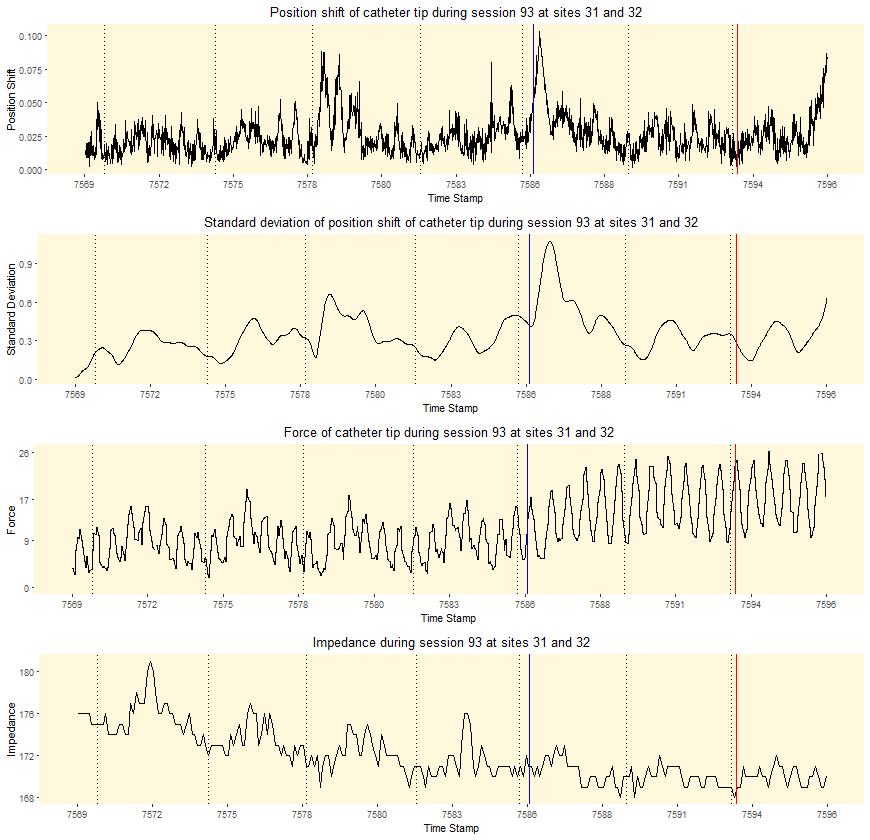

Supplementary figure 2: Reconstituted curves from VISITAG™ Module data export demonstrating first-annotated site “end” time-point at 16.3s (ACCURESP™ “off”, blue line) and 23.7s (ACCURESP™ “on”, red line) following RF onset in case 2, right PV. Inter-ablation site transition occurred with pure R UE morphology present at both site 1 completion and site 2 onset (ACCURESP™ “off” ILD 4.7mm); clear changes in the catheter tip position shift and (position) standard deviation (SD) occur just before ACCURESP™ “off” annotation, with additional change in CF noted. All curves are drawn black, since CF was maintained ≥1g and the catheter movement was sufficiently rapid to ensure that all catheter tip location data was annotated to either site 1, or 2 (ACCURESP™ “off”). Note: “Session 93 at sites 31 and 32” represents the unique automatically generated VISITAG™ Module annotated identifiers for this site.

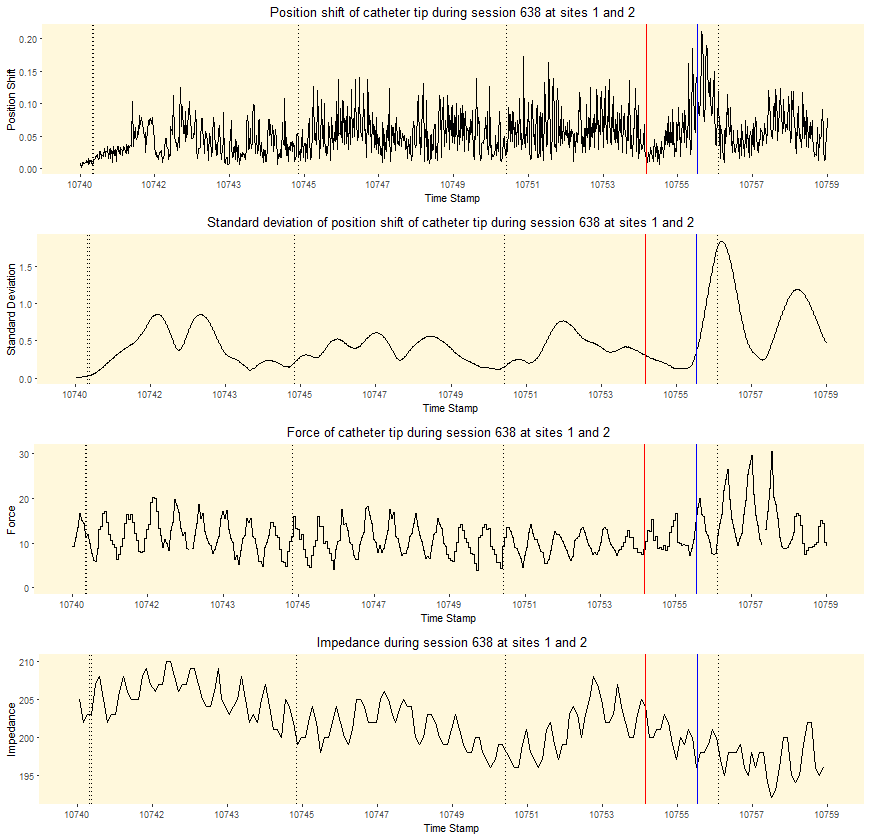

Supplementary figure 3: Reconstituted curves from VISITAG™ Module data export demonstrating first-annotated site “end” time-point at 14.4s (ACCURESP™ “on”, red line) and 15.7s (ACCURESP™ “off”, blue line) following RF onset in case 10, left PV. Inter-ablation site transition occurred with pure R UE morphology present at both site 1 completion and site 2 onset (ACCURESP™ “off” ILD 5.3mm); clear changes in the catheter tip position shift and (position) standard deviation (SD) occur concurrent with ACCURESP™ “off” annotation, with additional change in CF noted. All curves are drawn black, since CF was maintained ≥1g and the catheter movement was sufficiently rapid to ensure that all catheter tip location data was annotated to either site 1, or 2. Note: “Session 638 at sites 1 and 2” represents the unique automatically generated VISITAG™ Module annotated identifiers for this site.

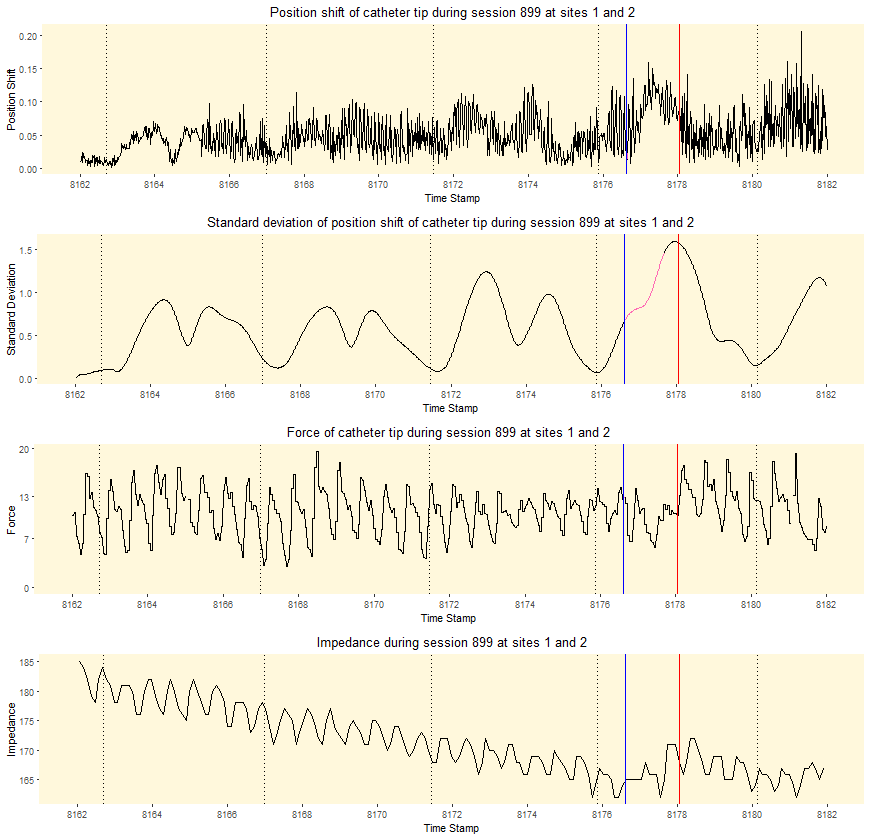

Supplementary figure 4: Reconstituted curves from VISITAG™ Module data export demonstrating first-annotated site “end” time-point at 14.5s (ACCURESP™ “off”, blue line) and 15.9s (ACCURESP™ “on”, red line) following RF onset in case 14, left PV. Inter-ablation site transition occurred with pure R UE morphology present at both site 1 completion and site 2 onset (ACCURESP™ “off” ILD 5.1mm); the peak position shift is noted mid-way between these annotation events, concurrent with subtle changes in CF and impedance. With ACCURESP™ “off” there was 0.05s non-annotated inter-ablation site time due to position instability >2*(2mm SD) – difficult to see on any curve aside the (position) standard deviation, where an additional 1s (i.e. 60 positions) is seen in view of the CARTO®3 SD calculation “system logic”. Note: “Session 899 at sites 1 and 2” represents the unique automatically generated VISITAG™ Module annotated identifiers for this site.

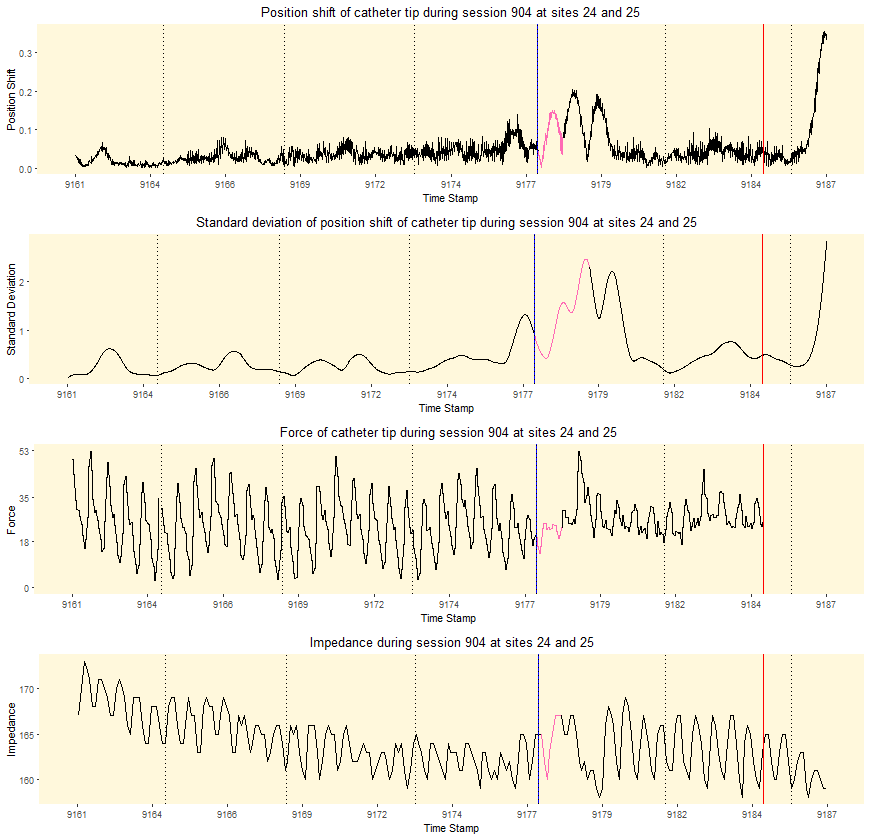

Supplementary figure 5: Reconstituted curves from VISITAG™ Module data export demonstrating first-annotated site “end” time-point at 15.9s (ACCURESP™ “off”, blue line) and 23.6s (ACCURESP™ “on”, red line) following RF onset in case 14, right PV. Inter-ablation site transition occurred with pure R UE morphology present at both site 1 completion and site 2 onset (ACCURESP™ “off” ILD 4.9mm); clear changes in all data curves occur concurrent with ACCURESP™ “off” annotation timing. With ACCURESP™ “off” there was 0.89s non-annotated inter-ablation site time due to position instability >2*(2mm SD); data curves return to black once all filter preferences have been met (a total of 1.89s pink is seen for position SD in view of the CARTO®3 SD calculation “system logic”). Note: “Session 904 at sites 24 to 25” represents the unique automatically generated VISITAG™ Module annotated identifiers for this site.

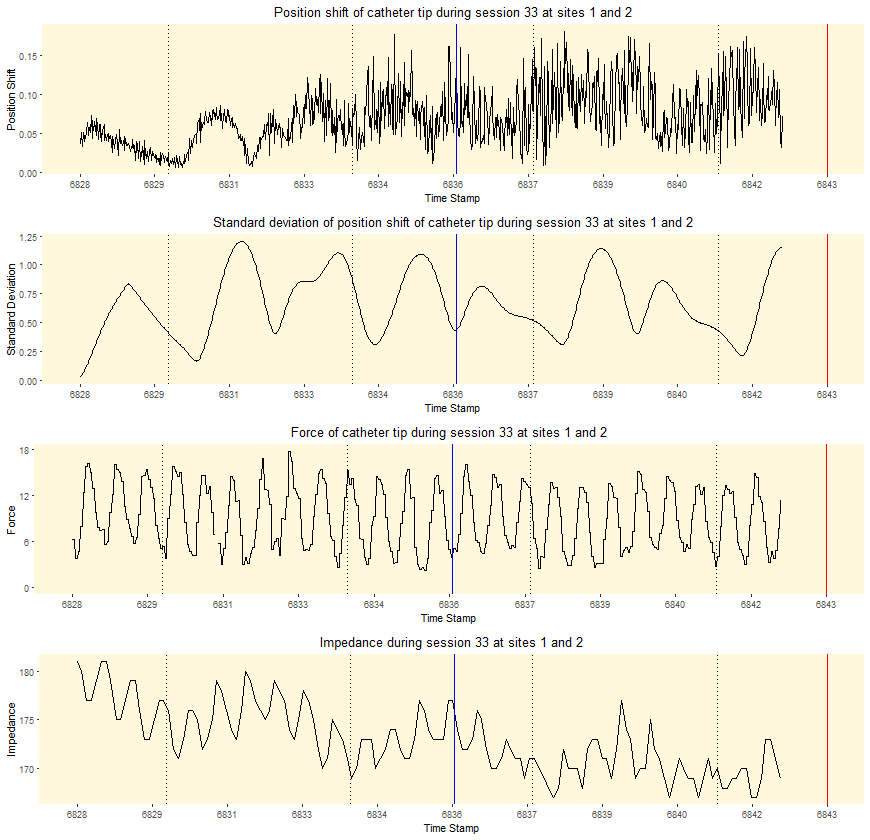

Supplementary figure 6: Reconstituted curves from VISITAG™ Module data export demonstrating first-annotated site “end” time-point at 7.9s (ACCURESP™ “off”, blue line, inadvertent movement) and 15.6s (ACCURESP™ “on”, red line) following RF onset in case 16, left PV. Inter-ablation site transition occurred with pure R UE morphology present at both site 1 completion and site 2 onset (ACCURESP™ “off” ILD 2.1mm); there are no clear changes in the plotted data. The curves end at 14.8s following RF onset since this time was concurrent with ACCURESP™ “off” annotation transition to LAPW site 3. Note: “Session 33 at sites 1 and 2” represents the unique automatically generated VISITAG™ Module annotated identifiers for this site.

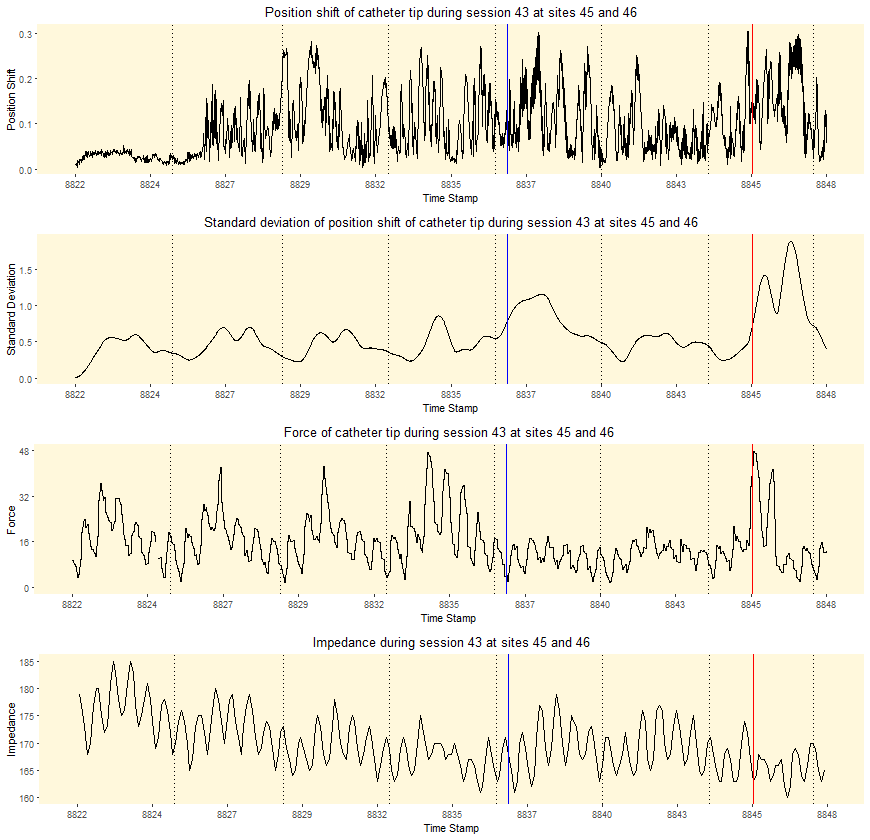

Supplementary figure 7: Reconstituted curves from VISITAG™ Module data export demonstrating first-annotated site “end” time-point at 15.2s (ACCURESP™ “off”, blue line) and 23.8s (ACCURESP™ “on”, red line) following RF onset in case 16, right PV. Inter-ablation site transition occurred with pure R UE morphology present at both site 1 completion and site 2 onset (ACCURESP™ “off” ILD 4.3mm); the peak position SD occurs concurrent with ACCURESP™ “off” annotation timing, with additional changes in CF and impedance evident at this time. All curves are drawn black, since CF was maintained ≥1g and the catheter movement was sufficiently rapid to ensure that all catheter tip location data was annotated to either site 1, or 2. Note: “Session 43 at sites 45 and 46” represents the unique automatically generated VISITAG™ Module annotated identifiers for this site.

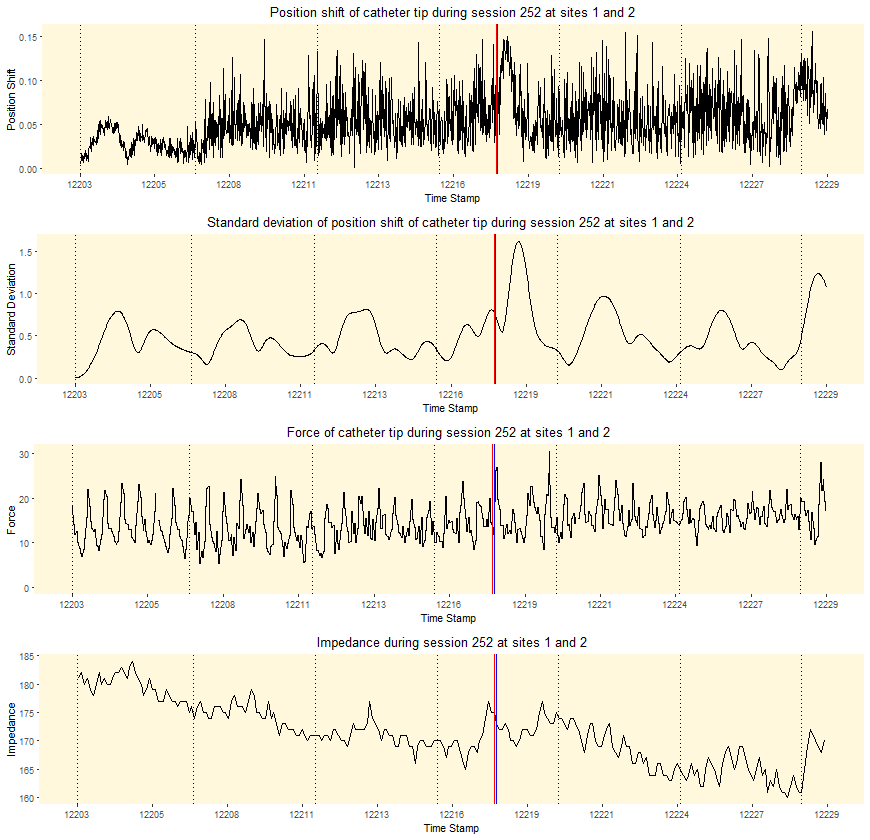

Supplementary figure 8: Reconstituted curves from VISITAG™ Module data export demonstrating first-annotated site “end” time-point at 14.8s (ACCURESP™ “on”, red line) and 14.9s (ACCURESP™ “off”, blue line) following RF onset in case 22, left PV. Inter-ablation site transition occurred with pure R UE morphology present at both site 1 completion and site 2 onset (ACCURESP™ “off” ILD 6.6mm); the peak position shift and SD occur just after this time-point; additional subtle increase in impedance is evident at this time. All curves are drawn black, since CF was maintained ≥1g and the catheter movement was sufficiently rapid to ensure that all catheter tip location data was annotated to either site 1, or 2. Note: “Session 252 at sites 1 and 2” represents the unique automatically generated VISITAG™ Module annotated identifiers for this site.

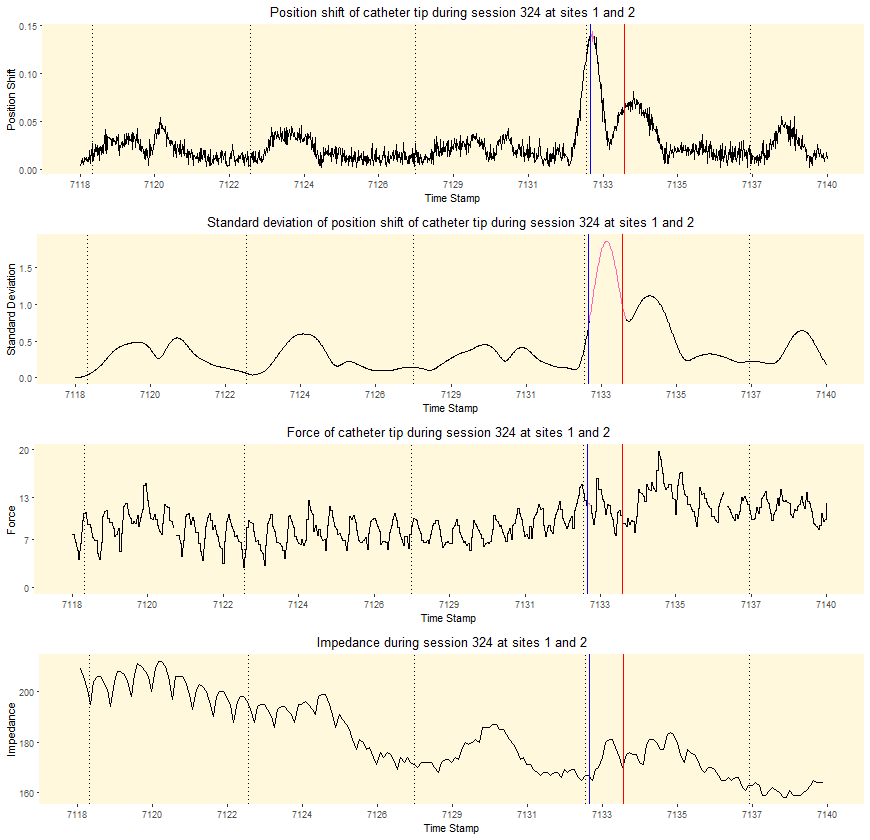

Supplementary figure 9: Reconstituted curves from VISITAG™ Module data export demonstrating first-annotated site “end” time-point at 15.0s (ACCURESP™ “off”, blue line) and 16.0s (ACCURESP™ “on”, red line) following RF onset in case 23, left PV. Inter-ablation site transition occurred with pure R UE morphology present at both site 1 completion and site 2 onset (ACCURESP™ “off” ILD 7.0mm); peak position shift coincides ACCURESP™ “off” annotation timing and a subtle increase in impedance. With ACCURESP™ “off” there was 0.12s non-annotated inter-ablation site time due to position instability >2*(2mm SD); data curves return to black once all filter preferences have been met (a total of 1.12s pink is seen for position SD in view of the CARTO®3 SD calculation “system logic”). Note: “Session 324 at sites 1 and 2” represents the unique automatically generated VISITAG™ Module annotated identifiers for this site.

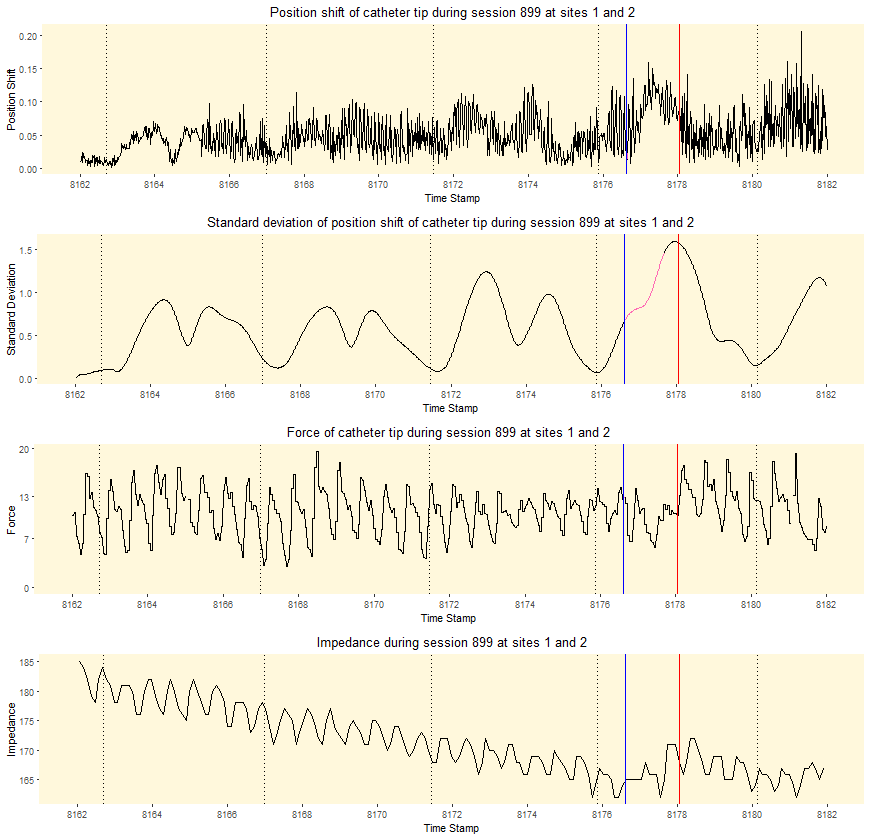

Supplementary figure 10: Reconstituted curves from VISITAG™ Module data export demonstrating first-annotated site “end” time-point at 14.5s (ACCURESP™ “off”, blue line) and 15.9s (ACCURESP™ “on”, red line) following RF onset in case 14, left PV. Inter-ablation site transition occurred with pure R UE morphology present at both site 1 completion and site 2 onset (ACCURESP™ “off” ILD 5.1mm); the peak position shift is noted mid-way between these annotation events, concurrent with subtle changes in CF and impedance. With ACCURESP™ “off” there was 0.05s non-annotated inter-ablation site time due to position instability >2*(2mm SD) – difficult to see on any curve aside the (position) standard deviation, where an additional 1s (i.e. 60 positions) is seen in view of the CARTO®3 SD calculation “system logic”. Note: “Session 899 at sites 1 and 2” represents the unique automatically generated VISITAG™ Module annotated identifiers for this site. This figure represents duplication of supplementary figure 4, but is shown here in series with supplementary figures 11-13 to permit greater ease of review of annotation timing changes during continuous LAPW RF application in this single case.

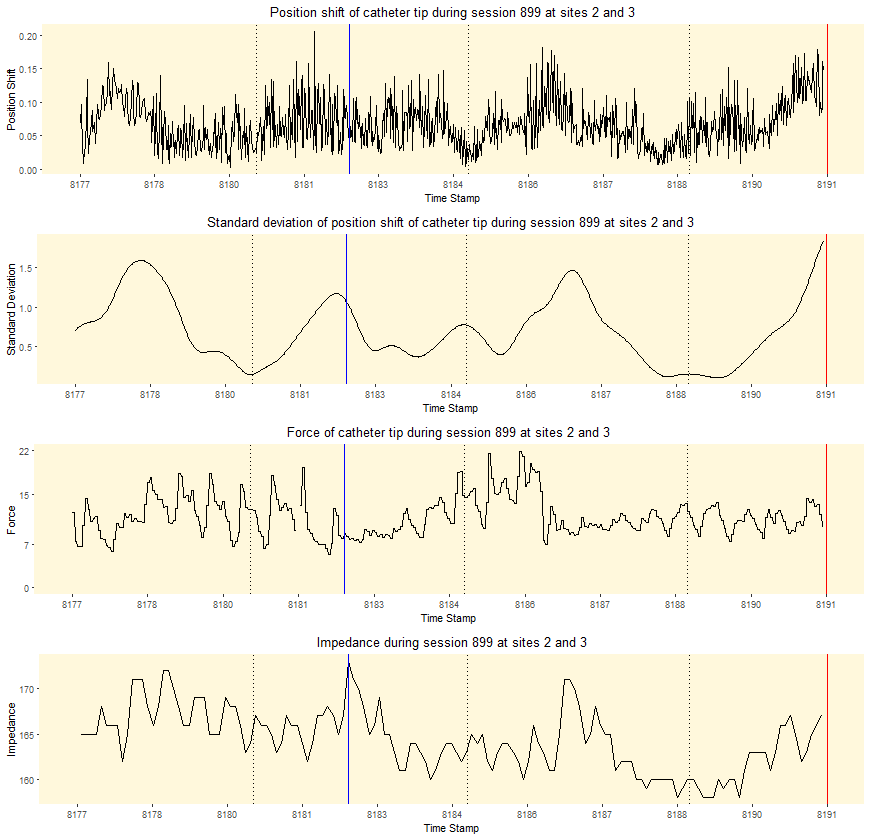

Supplementary figure 11: Reconstituted curves from VISITAG™ Module data export demonstrating second-annotated site “end” time-point following an additional 5.3s RF with ACCURESP™ “off” (blue line) and 13.2s with ACCURESP™ “on” (red line) during continuous LAPW RF in case 14, left PV. Inter-ablation site transition occurred with pure R UE morphology present at both site 2 completion and site 3 onset (ACCURESP™ “off” ILD 2.4mm); the peak position shift is noted just before ACCURESP™ “off” annotation, concurrent with subtle changes in CF and impedance. All curves are drawn black, since CF was maintained ≥1g and the catheter movement was sufficiently rapid to ensure that all catheter tip location data was annotated to either site 2, or 3. Note: “Session 899 at sites 2 and 3” represents the unique automatically generated VISITAG™ Module annotated identifiers for this site.

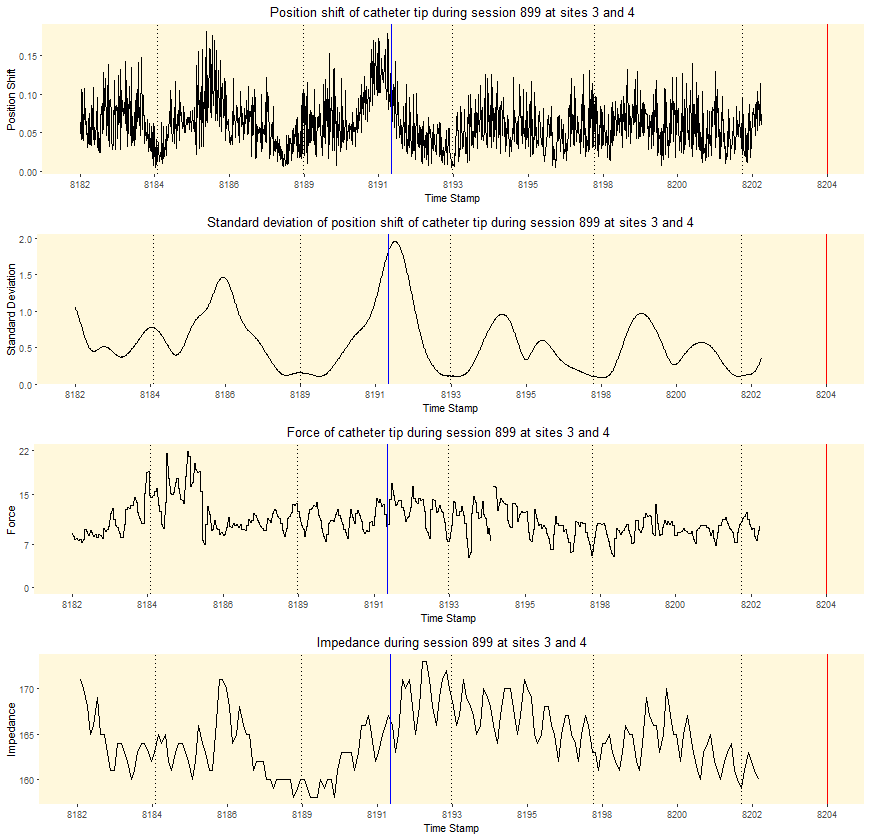

Supplementary figure 12: Reconstituted curves from VISITAG™ Module data export demonstrating third-annotated site “end” time-point following an additional 9.3s RF with ACCURESP™ “off” (blue line) and 12.8s with ACCURESP™ “on” (red line) during continuous LAPW RF in case 14, left PV. Transition to site 4 with ACCURESP “off” coincided with UE morphology change from pure R at site 3 completion, to RS at site 4 onset (ACCURESP™ “off” ILD 4.8mm, time to pure R UE morphology change at site 4, 1.3s); the peak position shift and position SD coincide with ACCURESP™ “off” annotation, concurrent with a clear change in impedance. All curves are drawn black, since CF was maintained ≥1g and the catheter movement was sufficiently rapid to ensure that all catheter tip location data was annotated to either site 3, or 4. The curves end at 40.1s following RF onset since this time was concurrent with ACCURESP™ “off” annotation transition to LAPW site 5. Note: “Session 899 at sites 3 and 4” represents the unique automatically generated VISITAG™ Module annotated identifiers for this site.

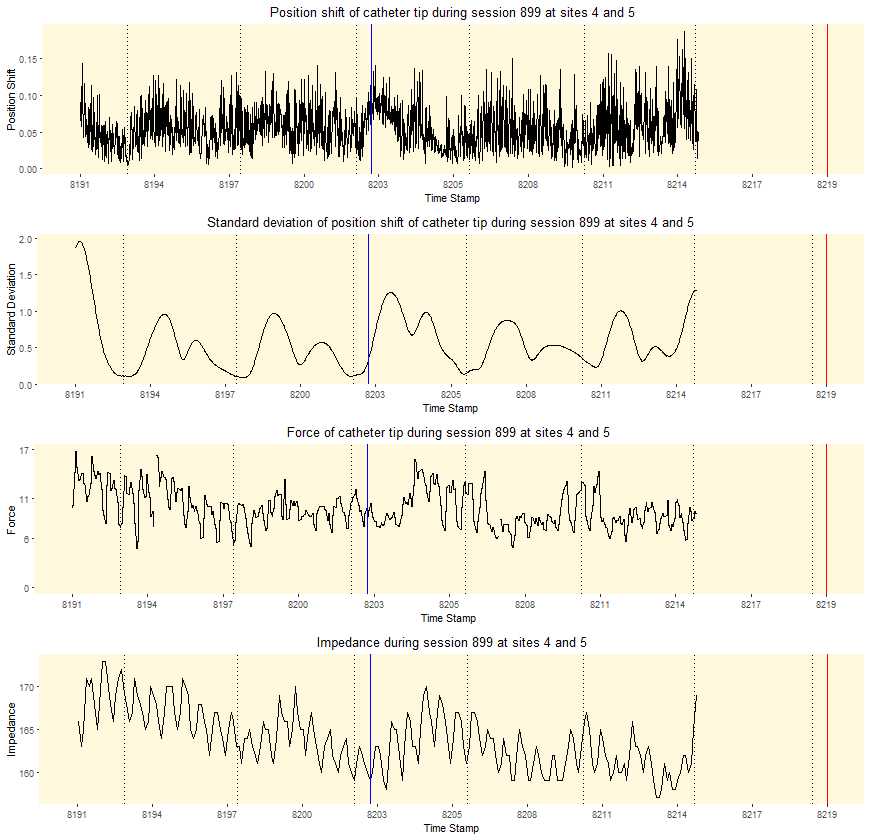

Supplementary figure 13: Reconstituted curves from VISITAG™ Module data export demonstrating fourth-annotated site “end” time-point following an additional 11.0s RF with ACCURESP™ “off” (blue line) and 15.1s with ACCURESP™ “on” (red line) during continuous LAPW RF in case 14, left PV. Transition to site 5 with ACCURESP “off” occurred with pure R UE morphology present at both site 4 completion and site 5 onset (ACCURESP™ “off” ILD 5.4mm); the peak position SD occurs just after ACCURESP™ “off” annotation, concurrent with a subtle change in CF and impedance. All curves are drawn black, since CF was maintained ≥1g and the catheter movement was sufficiently rapid to ensure that all catheter tip location data was annotated to either site 4, or 5. The curves end at 52.4s following RF onset since this time was concurrent with ACCURESP™ “off” annotation transition to LAPW site 5. Note: “Session 899 at sites 4 and 5” represents the unique automatically generated VISITAG™ Module annotated identifiers for this site.

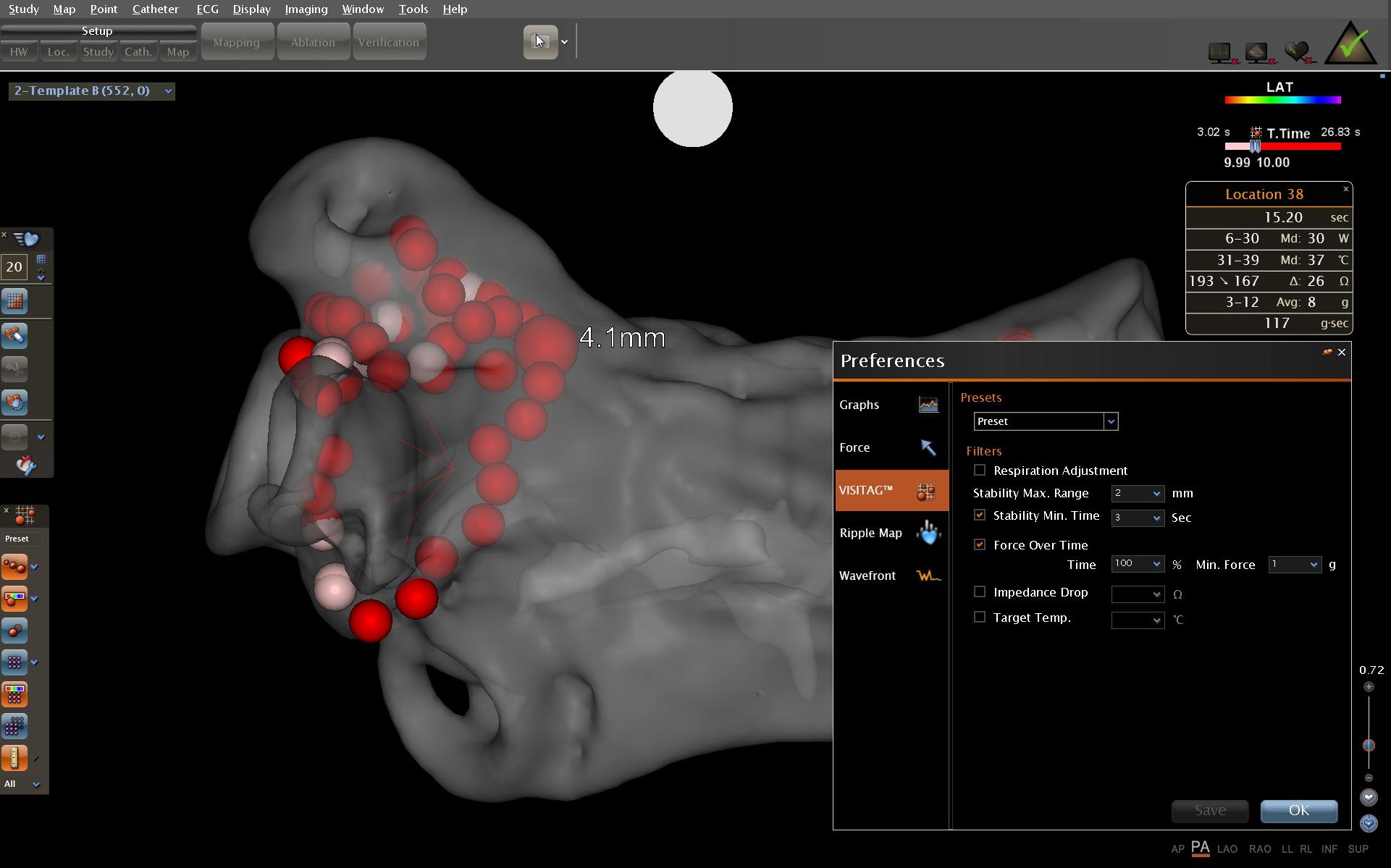

A

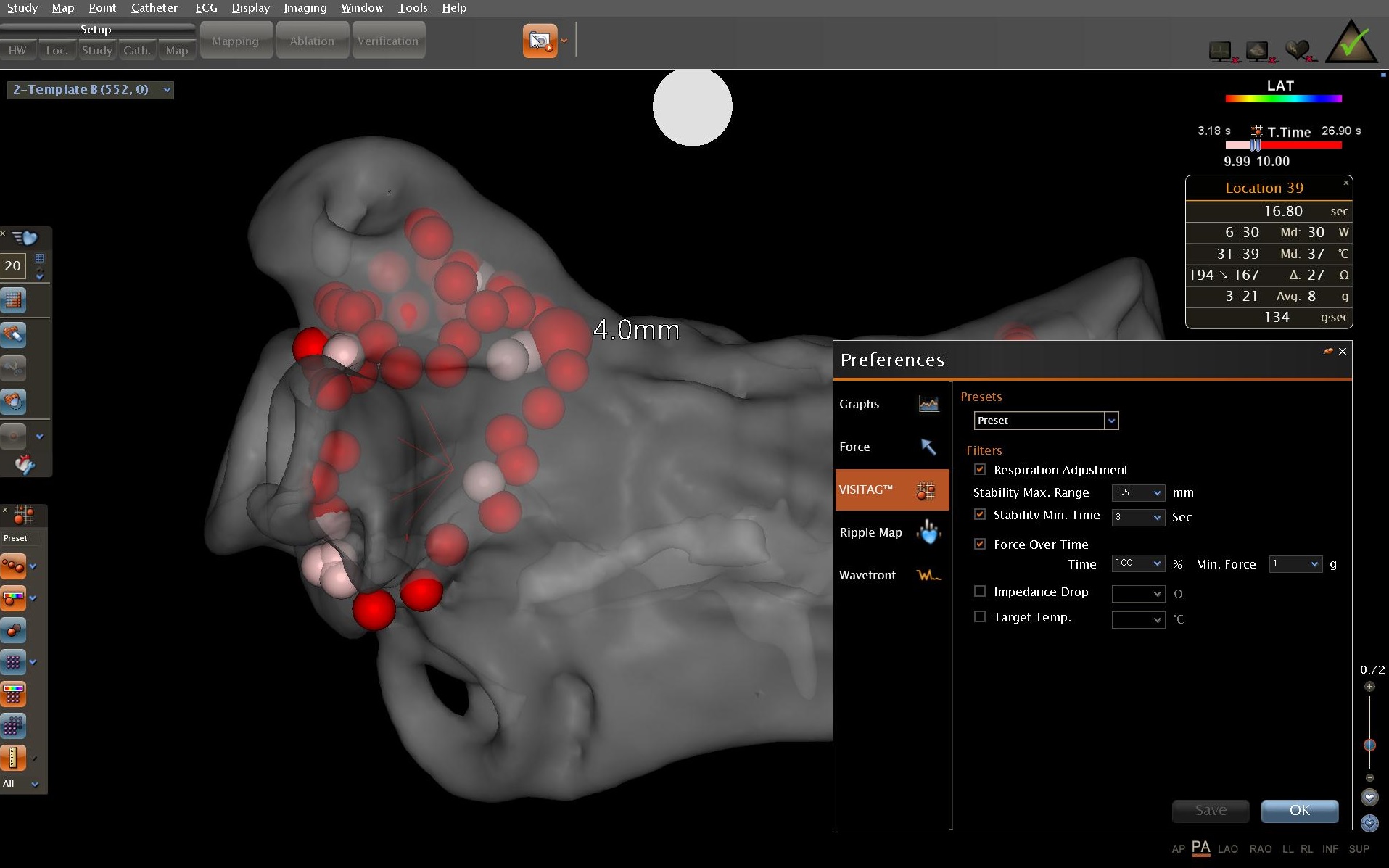

B

Supplementary figure 14: Case 5 (left PV, PA view) annotated site 1 data with VISITAG™ Module preference settings shown; (A) ACCURESP™ “off” at 2mm position stability, and (B) ACCURESP™ “on” at 1.5mm position stability. Annotated parameters are shown as “Location 38” and “Location 39”, respectively and indicate a 1.6s delay to annotated transition to site 2, using ACCURESP “on”. Site 1-2 ILD also shown – 4.1mm and 4.0mm.
